# TGF-β- and IL-1-dependent fibroblast states differentially shape pancreatic cancer microenvironments and metastasis

**DOI:** 10.64898/2026.09.08.750042

**Authors:** Priscilla S.W. Cheng, Eloise G. Lloyd, Muntadher Jihad, Marta Zaccaria, Kristen Burgess, Gianluca Mucciolo, Sneha Harish, Weike Luo, Judhell S. Manansala, Wenlong Li, Joaquín Araos Henríquez, Debasmita Mukherjee, Sally Mills, Paul M. Johnson, Alejandro Alonso Montero, Melinda J. Duer, Paul D.W. Kirk, Mireia Vallespinos, Giulia Biffi

## Abstract

Pancreatic ductal adenocarcinoma (PDAC) contains heterogeneous cancer-associated fibroblast (CAF) populations, including interleukin 1 (IL-1)-dependent inflammatory CAFs (iCAFs) and transforming growth factor beta (TGF-β)-dependent myofibroblastic CAFs (myCAFs), whose functions at primary and metastatic sites remain incompletely defined. Here, we genetically disrupted IL-1 or TGF-β signalling in fibroblast activation protein (FAP)-expressing CAFs arising from a shared cellular origin to determine how myCAF and iCAF states shape PDAC progression and microenvironments. Depletion of TGF-β-dependent myCAFs, but not IL-1-dependent iCAFs, did not alter primary tumour growth or local metastatic dissemination but significantly reduced liver and lung metastases. Matched primary tumours and liver metastasis analyses also revealed site-specific stromal responses to TGF-β signalling disruption. Experimental models of liver metastasis colonisation and outgrowth indicated that the reduced metastatic phenotype was linked primarily to alterations within pancreatic tumours rather than to impaired malignant cell outgrowth at metastatic sites. Single-cell RNA-sequencing of myCAF-depleted primary tumours revealed extensive transcriptional reprogramming across fibroblast, immune and malignant compartments, highlighting the complexity of assigning myCAF-to-malignant cell signalling mechanisms *in vivo*. To resolve this crosstalk, we co-cultured PDAC organoids with pancreatic stellate cells engineered to remain in myCAF or iCAF states. Cross-model transcriptomic analyses, together with genetic and pharmacological perturbation studies, showed that myCAF-derived TGF-β directly promotes epithelial-to-mesenchymal transition signalling in PDAC malignant cells, providing a candidate mechanism for the reduced metastasis observed upon myCAF depletion *in vivo*. These findings reveal distinct, context-dependent roles for TGF-β- and IL-1-dependent CAF states and identify myCAF-derived TGF-β signalling as a stromal mechanism promoting malignant cell plasticity.

**STATEMENT OF SIGNIFICANCE:** Manipulation of inflammatory fibroblast and myofibroblast states shows that distinct fibroblast programmes arising from a shared cellular origin differentially shape pancreatic cancer progression and context-dependent metastasis formation.

## INTRODUCTION

Pancreatic Ductal Adenocarcinoma (PDAC) is characterised by aggressive local growth and early metastatic dissemination, contributing to dismal patient outcomes^1^. A contributor to these challenges is the extensive fibroblast-rich tumour microenvironment (TME), whose functional complexity remains incompletely understood. Three major cancer-associated fibroblast (CAF) states have been described in murine and human PDAC: abundant myofibroblastic and inflammatory CAFs (myCAFs and iCAFs), and a smaller subset of major histocompatibility complex class II (MHC)^+^ antigen-presenting CAFs (apCAFs)^2–7^. Within and across these states, additional subsets have been identified, highlighting the remarkable phenotypic diversity of PDAC CAFs^8,9^. This heterogeneity is driven by multiple factors, including variable cues from malignant cells and distinct cells of origin^5,8–16^. For example, while apCAFs predominantly arise from mesothelial cells, both iCAFs and myCAFs can arise from pancreatic stellate cells (PSCs) in response to malignant cell-derived interleukin-1 (IL-1) and transforming growth factor beta (TGF-β), respectively^2,17^. Importantly, CAF states are not fixed, but can change according to spatial cues, cytokine gradients and disease stage^6,7,18,19^.

Although myCAF and iCAF states are abundant in PDAC and other solid tumours^7,20^, their *in vivo* functions in primary and metastatic tumours remain incompletely understood. This knowledge gap has hindered the rational development of therapeutic strategies that could systemically target similarly activated CAF states across primary and metastatic sites. Early attempts to target the PDAC stroma, including global fibroblast depletion or Hedgehog pathway inhibition, have yielded disappointing outcomes, underscoring the need for a more nuanced understanding of CAF states and function^21–23^. Indeed, both tumour-promoting and tumour-restraining roles have been described for myCAFs. Previous studies suggest that depletion of alpha smooth muscle actin-positive (αSMA^+^) myCAFs, or their product collagen I, can accelerate PDAC progression in mouse models, suggesting myCAFs and extracellular matrix (ECM) components can restrain tumour growth^24,25^. In contrast, we and others have identified myCAF subsets that can promote PDAC progression and immunosuppression^8,14^. Thus, distinct subsets of CAFs appear to have diverse functions that exert opposing effects on tumour progression. While genetic manipulation of pancreatic CAFs within the primary TME has local stromal effects^26^, no study has independently depleted myCAF and iCAF states arising from a shared cellular origin to define their *in vivo* functions at primary and metastatic sites.

To address this knowledge gap, we developed genetically engineered mouse models (GEMMs) in which myCAF- or iCAF-associated signalling can be disrupted in fibroblast activation protein (FAP)-expressing stromal cells across organs. In parallel, we engineered PSC lines to remain ‘locked’ in either myCAF or iCAF states^27^. Together, these approaches enable interrogation of state-specific PDAC CAF functions across disease sites.

## RESULTS

### Characterisation of novel GEMMs for depletion of *Fap*-expressing TGF-β- or IL-1-dependent CAFs in PDAC

Because IL-1 and TGF-β signalling promote iCAF and myCAF phenotypes, respectively, we targeted these pathways to perturb their associated CAF states in PDAC. To restrict these perturbations to CAFs arising from a shared mesenchymal origin, we targeted *Fap*-expressing cells. First, human PDAC iCAFs and myCAFs express FAP^3,28^. Second, FAP expression is largely restricted to fibroblast populations rather than other stromal cell types in adult mice^29,30^. Third, FAP^+^ CAFs have been implicated in diverse aspects of PDAC biology^15,31^. Fourth, *Fap*^High^ fibroblasts in pancreatitis represent putative precursors of PDAC CAFs^32^. Targeting *Fap*-expressing cells also minimised perturbations of apCAFs, which can also respond to TGF-β and IL-1 signalling but arise predominantly from FAP^−^ mesothelial cells^17^. Indeed, analysis of single-cell RNA-sequencing (scRNA-seq) profiles from tumours derived from orthotopic transplantation of PDAC organoids in mice confirmed that *Fap* expression was localised predominantly in fibroblasts, was present in >50% of iCAFs and myCAFs, and was low in apCAFs (**Supplementary Fig. 1A**).

To ablate TGF-β signalling in *Fap*-expressing cells, we crossed previously described *Fap*-tTA and Tet-O-Cre mice^30^ with *Tgfbr2^fl/fl^*^33^ and *Rosa26^LSL-tdTom/tdTom^* mice, generating *Fap*-tTA; Tet-O-Cre; *Rosa26^LSL-tdtom/tdtom^*; *Tgfbr2^fl/fl^* mice (hereafter referred to as FTC; **Supplementary Fig. 1B**). In this model, doxycycline (dox) withdrawal induces Cre-mediated TdTomato expression and *Tgfbr2* deletion in *Fap*-expressing cells. FTC mice maintained on dox, as well as on dox or off dox *Fap*-tTA; *Rosa26^LSL-tdtom/tdtom^*; *Tgfbr2^fl/fl^* mice lacking Tet-O-Cre (hereafter, FT), were used as controls. Six weeks following dox withdrawal, TdTomato expression was detected in the pancreas of FTC mice, confirming Cre activity, whereas TdTomato signal was absent in FT controls and significantly lower in FTC controls on dox (**Supplementary Fig. 1C-D**). Orthotopic transplantation of *Kras^LSL-G12D/+^*; *Trp53^LSL-R^*^172*H*/+^; *Pdx1*-Cre (KPC) GEMM-derived PDAC organoids^8,34–36^ into FT and FTC mice after six weeks of dox withdrawal, or into FT and FTC mice maintained on dox, was performed (**Fig. 1A**). Analysis of PDAC tumours by immunohistochemistry revealed TdTomato expression, marking cells targeted via *Fap*-driven Cre recombination, in FTC tumours following dox withdrawal (**Fig. 1B**). Flow cytometry analysis of tumours demonstrated that ∼30% of podoplanin (PDPN)^+^ fibroblasts were TdTomato^High^, whereas TdTomato high-positivity was detected in ∼5% of endothelial cells, potentially undergoing endothelial-to-mesenchymal transition^37^, and < 1% of immune or epithelial cells (**Fig. 1C** and **Supplementary Fig. 1E**). Given that *Fap* is expressed by a subset of pericytes (**Supplementary Fig. 1A**), we further assessed TdTomato expression in this compartment. Analysis of a murine PDAC scRNA-seq dataset identified pericytes as platelet-derived growth factor receptor alpha-negative (*Pdgfra*^-^), *Pdpn*^-^, *Acta2*^+^ cells, with *Acta2* encoding αSMA, whereas fibroblasts expressed these markers to varying degrees (**Supplementary Fig. 1F**). Multiplex immunofluorescence (IF) analysis revealed TdTomato expression in ∼10% of PDGFRα^-^ PDPN^-^ αSMA^+^ pericytes and in ∼20% of PDPN^+^ fibroblasts (**Fig. 1D** and **Supplementary Fig. 1G**). Fibroblasts comprised ∼20% of cells within primary TMEs from both FT and FTC mice, whereas pericytes accounted for only ∼2% (**Supplementary Fig. 1H**), indicating that fibroblasts represent the major stromal compartment in which TGF-β signalling is disrupted. Notably, given that pericytes can contribute to CAF composition^7,38,39^, these models primarily target FAP^+^ fibroblasts but may also capture contributions from other FAP^+^ stromal populations. To assess recombination efficiency at the genomic level, TdTomato^High^ and TdTomato^−^ PDPN^+^ fibroblasts, together with EpCAM^+^ PDPN^−^ epithelial cells largely derived from transplanted PDAC organoids, were flow-sorted from tumours. Quantitative analysis revealed > 90% deletion of the *Tgfbr2* locus in TdTomato^High^ fibroblasts from FTC tumours following dox withdrawal, whereas no recombination was detected in fibroblasts from FT and FTC ON dox control tumours (**Supplementary Fig. 1I-J**). As expected, only the wild-type *Tgfbr2* allele was detected in epithelial cells (**Supplementary Fig. 1J**).

**Figure 1.**
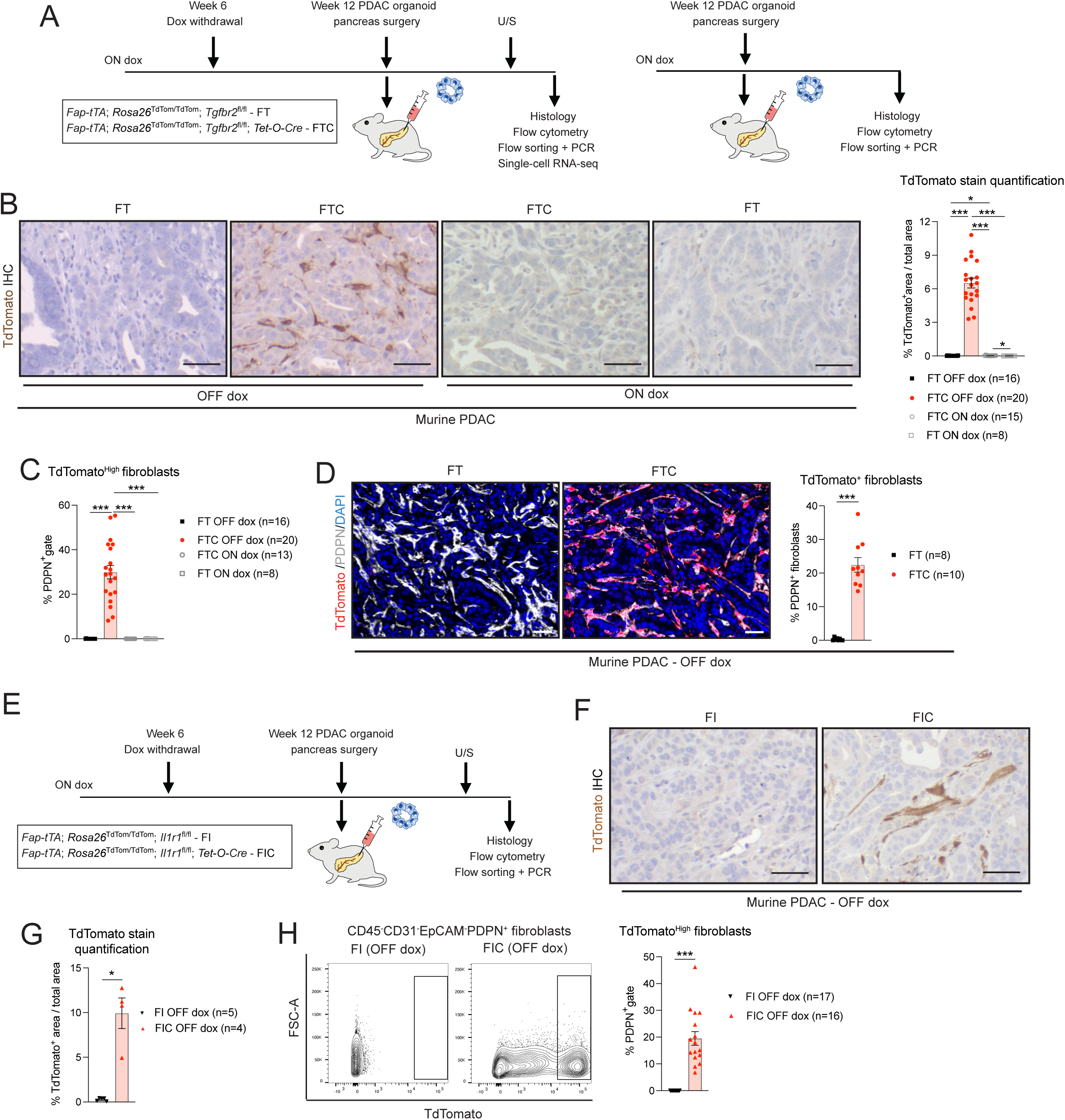
Characterisation of novel mouse models for depletion of *Fap*-expressing TGF-β-or IL-1-dependent fibroblasts in PDAC. **(A)** Schematic of experimental design of analyses of orthotopic transplantation models of pancreatic ductal adenocarcinoma (PDAC) organoids in *Fap-tTa; Rosa26^LSL-tdTom/tdTom^; Tgfbr2^fl/fl^* (hereafter, FT) and *Fap-tTa; tet-O-Cre; Rosa26^LSL-tdTom/tdTom^; Tgfbr2^fl/fl^* (hereafter, FTC) mice with or without doxycycline (dox) withdrawal. U/S, ultrasound. **(B)** Left: Representative TdTomato immunohistochemistry (IHC) stains in FT and FTC PDAC after dox withdrawal or in FT and FTC PDAC on dox. Scale bars, 25 μm. Right: Quantification of TdTomato stains in FT (n=16) and FTC (n=20) PDAC after dox withdrawal or in FT (n=8) and FTC (n=15) PDAC on dox. Results show mean ± standard error of the mean (SEM). * *P adj* < 0.05; *** *P adj* < 0.001, Kruskal-Wallis test. **(C)** Flow cytometric analyses of TdTomato^High^ CAFs (CD45^-^CD31^-^EpCAM^-^ PDPN^+^) from the parental PDPN^+^ gate in FT (n=16) and FTC (n=20) PDAC after dox withdrawal or in FT (n=8) and FTC (n=13) PDAC on dox. Results show mean ± SEM. *** *P adj* < 0.001, Kruskal-Wallis test. **(D)** Left: Representative multiplex immunofluorescence (mIF) images of TdTomato (red), PDPN (white) and DAPI (nuclear stain, blue) in FT and FTC PDAC tissues after dox withdrawal. Scale bars, 50 μm. Right: Quantification of TdTomato^+^PDPN^+^ECAD^-^DAPI^+^ cells relative to PDPN^+^ECAD^-^DAPI^+^ fibroblasts in FT (n=8) and FTC (n=10) PDAC tissues after dox withdrawal. Results show mean ± SEM. *** *P* < 0.001, Mann-Whitney test. **(E)** Schematic of experimental design of analyses of orthotopic transplantation models of PDAC organoids in *Fap-tTa; Rosa26^LSL-tdTom/tdTom^; Il1r1^fl/fl^* (hereafter, FI) and *Fap-tTa; tet-O-Cre; Rosa26^LSL-tdTom/tdTom^; Il1r1^fl/fl^* (hereafter, FIC) mice with dox withdrawal. **(F)** Representative TdTomato IHC stains in FI and FIC PDAC after dox withdrawal. Scale bars, 25 μm. **(G)** Quantification of TdTomato stains in FI (n=5) and FIC (n=4) PDAC after dox withdrawal. Results show mean ± SEM. * *P* < 0.05, Mann-Whitney test. **(H)** Left: Representative flow plots of TdTomato^High^ CAFs (CD45^-^CD31^-^EpCAM^-^PDPN^+^) from FI and FIC tumours after dox withdrawal. Right: Flow cytometric analyses of TdTomato^High^ CAFs from the parental PDPN^+^ gate in FI (n=17) and FIC (n=16) PDAC after dox withdrawal. Results show mean ± SEM. *** *P* < 0.001, Mann-Whitney test.

To ablate IL-1 signalling in *Fap*-expressing fibroblasts, we crossed *Fap*-tTA and Tet-O-Cre mice^30^ with *Il1r1^fl/fl^* mice^40^ and *Rosa26^LSL-tdTom/tdTom^* mice, generating *Fap*-tTA; Tet-O-Cre; *Rosa26^LSL-tdtom/tdtom^*; *Il1r1^fl/fl^* mice (hereafter, FIC; **Supplementary Fig. 1K**). In FIC mice, dox withdrawal induces Cre-mediated TdTomato expression and *Il1r1* deletion in *Fap*-expressing cells. *Fap*-tTA; *Rosa26^LSL-tdtom/tdtom^*; *Il1r1^fl/fl^* mice lacking Tet-O-Cre (hereafter, FI), were used as controls. Orthotopic transplantation of PDAC organoids into FI and FIC mice was also performed after six weeks of dox withdrawal (**Fig. 1E**). Analysis of PDAC tumours by immunohistochemistry revealed TdTomato expression in FIC tumours following dox withdrawal, and flow cytometry confirmed TdTomato positivity in ∼20% of PDPN^+^ fibroblasts (**Fig. 1F-H**). In this model, recombination efficiency was assessed at the genomic level using TdTomato^+^, rather than TdTomato^High^, fibroblasts, together with TdTomato^−^ fibroblasts and epithelial cells flow-sorted from tumours. Potentially reflecting the inclusion of TdTomato^Low^ fibroblasts, which may represent a heterogeneous population with variable knockout efficiency, quantitative analysis revealed an average of 50% deletion of the *Il1r1* locus in TdTomato^+^ fibroblasts from FIC tumours following dox withdrawal, whereas no recombination was detected in fibroblasts from FI control tumours (**Supplementary Fig. 1L**).

Together, these data establish genetically controlled models that efficiently disrupt TGF-β or IL-1 signalling in *Fap*-expressing CAFs in PDAC.

### Disruption of TGF-β or IL-1 signalling in *Fap*-expressing CAFs differentially remodels PDAC tumours

We first assessed the impact of disrupting TGF-β signalling in *Fap*-expressing CAFs on PDAC stromal composition by evaluating overall fibroblast abundance and collagen deposition. Flow cytometry showed comparable overall PDPN⁺ fibroblast abundance across groups, whereas Masson’s trichrome and picrosirius red staining revealed significantly reduced collagen accumulation in FTC tumours relative to FTC ON dox and FT controls (**Fig. 2A** and **Supplementary Fig. 2A-C**). Given the reduction in collagen accumulation, we next examined whether altered ECM composition was associated with changes in tumour mechanical properties. Rheological analysis^41^ showed that FTC tumours had significantly reduced storage and loss moduli compared with FT tumours, indicating a softer, less elastic and mechanically compromised matrix (**Fig. 2B** and **Supplementary Fig. 2D**). Together, these data indicate that TGFBR2 signalling in CAFs contributes to ECM deposition, stiffness and organisation in primary PDAC tumours.

**Figure 2.**
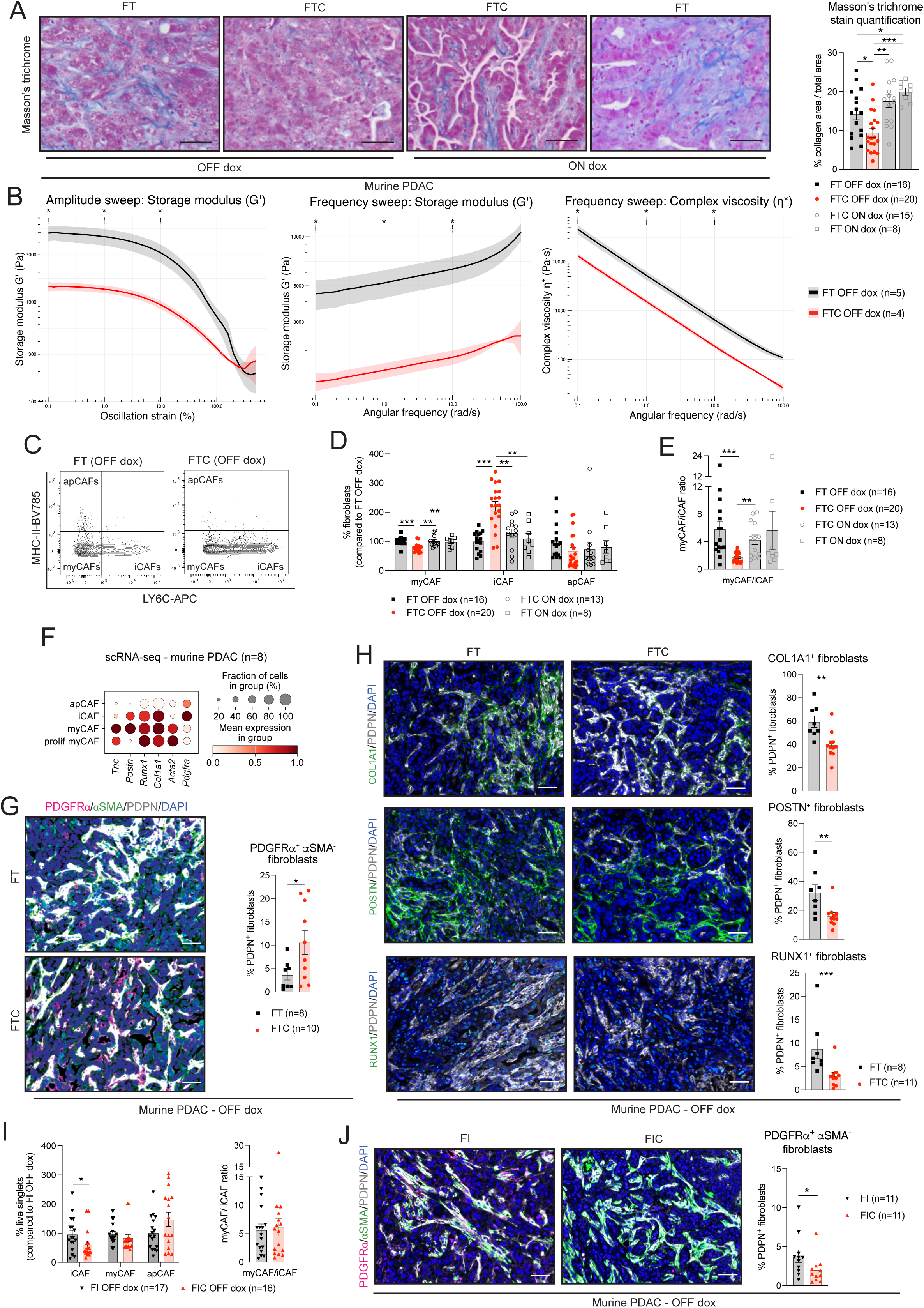
Disruption of TGF-β or IL-1 signalling in *Fap*-expressing fibroblasts differentially remodels PDAC tumours. **(A)** Left: Representative Masson’s trichrome stains in FT and FTC PDAC after dox withdrawal or on dox. Scale bars, 25 μm. Right: Quantification of Masson’s trichrome stain in FT (n=16) and FTC (n=20) PDAC after dox withdrawal or in FT (n=8) and FTC (n=15) PDAC on dox. Results show mean ± SEM. * *P adj* < 0.05; ** *P adj* < 0.01; *** *P adj* < 0.001; Kruskal-Wallis test. **(B)** Oscillatory rheological properties of *ex vivo* FT (n=5) and FTC (n=4) PDAC tumours after dox withdrawal under hydrated conditions at 37 °C: storage moduli from amplitude and frequency sweeps. Amplitude sweeps (left) were performed at a constant angular frequency at 1 rad/s, while frequency sweeps were performed at a constant strain of 0.5%. Complex viscosity (right) was calculated from frequency sweeps performed at a constant strain of 0.5%. **\*** *P* < 0.05, unpaired two-sided Welch’s *t*-test at 0.1%, 1% and 10% strain for amplitude sweeps and at 0.1, 1, and 10 rad/s for frequency sweeps. **(C)** Representative flow plots of myCAFs (Ly6C^-^MHCII^-^), iCAFs (Ly6C^+^MHCII^-^) and apCAFs (Ly6C^-^MHCII^+^) from FT and FTC PDACs after dox withdrawal. **(D-E)** Flow cytometric analyses of myCAFs, iCAFs and apCAFs from the parental PDPN^+^ gate **(D)** and myCAF/iCAF ratio **(E)** in FT (n=16) and FTC (n=20) PDAC after dox withdrawal or in FT (n=8) and FTC (n=13) PDAC on dox. Results show mean ± SEM. ** *P adj* < 0.01; *** *P adj* < 0.001; Kruskal-Wallis test. **(F)** Dot plot of scaled expression of *Pdgfra* and myofibroblastic markers in each CAF cluster of murine PDAC tissues (n=8), as analysed by single-cell RNA-sequencing (scRNA-seq). The colour intensity represents the expression level, and the size of the dots represents the percentage of expressing cells. Data are from Araos and Jihad et al^84^. **(G)** Left: Representative mIF images of αSMA (green), PDGFRα (pink), PDPN (white) and DAPI (blue) in FT and FTC PDAC tissues after dox withdrawal. Scale bars, 50 μm. Right: Quantification of PDGFRα^+^αSMA^-^PDPN^+^ECAD^-^DAPI^+^ fibroblasts relative to PDPN^+^ECAD^-^DAPI^+^ fibroblasts in FT (n=8) and FTC (n=10) PDAC tissues after dox withdrawal. Results show mean ± SEM. * *P* < 0.05, Mann-Whitney test. **(H)** Left: Representative mIF images of COL1A1/POSTN/RUNX1 (green), PDPN (white) and DAPI (blue) in FT and FTC PDAC tissues after dox withdrawal. Scale bars, 50 μm. Right: Quantification of COL1A1^+^ or POSTN^+^ or RUNX1^+^ PDPN^+^CK19^-^DAPI^+^ fibroblasts relative to PDPN^+^CK19^-^DAPI^+^ fibroblasts in FT (n=8) and FTC (n=11) PDAC tissues after dox withdrawal. Results show mean ± SEM. ** *P* < 0.01, *** *P* < 0.001, Mann-Whitney test. **(I)** Flow cytometric analyses of iCAFs, myCAFs, and apCAFs from live singlets (left) or myCAF/iCAF ratio (right) in FI (n=17) and FIC (n=16) PDAC after dox withdrawal. Results show mean ± SEM. * *P* < 0.05, Mann-Whitney test. **(J)** Left: Representative mIF images of αSMA (green), PDGFRα (pink), PDPN (white) and DAPI (blue) in FI and FIC PDAC tissues after dox withdrawal. Scale bars, 50 μm. Right: Quantification of PDGFRα^+^αSMA^-^PDPN^+^CK19^-^DAPI^+^ fibroblasts relative to PDPN^+^CK19^-^DAPI^+^ fibroblasts in FI (n=11) and FIC (n=11) PDAC tissues after dox withdrawal. Results show mean ± SEM. * *P* < 0.05, Mann-Whitney test.

Despite these stromal changes, the overall abundance of immune, endothelial, and epithelial cells was unchanged in FTC tumours compared to FT controls (**Supplementary Fig. 2E**). Similarly, flow cytometric analysis revealed no significant differences in the numbers of major immune cell populations, including macrophages, neutrophils, T cells, B cells and Natural Killer cells in FTC tumours compared to FT controls (**Supplementary Fig. 2F-G**). We next examined whether disruption of TGF-β signalling in CAFs altered fibroblast composition. Using a previously established flow cytometry strategy^8,42^, we observed a significant reduction in myCAFs (Ly6C^-^MHCII^-^) in FTC PDAC tumours relative to FTC ON dox and FT controls (**Fig. 2C-D**). In contrast, iCAFs (Ly6C^+^MHCII^-^) were increased, resulting in a significantly reduced myCAF/iCAF ratio, whereas apCAFs (MHCII^+^Ly6C^-^) were unchanged, consistent with their low *Fap* expression (**Fig. 2C-E**). Multiplex IF analysis for inflammatory and myofibroblastic CAF markers further supported a less myofibroblastic composition in FTC tumours, revealing an increased abundance of previously described PDGFRα^+^αSMA^-^ inflammatory fibroblasts^32^ and a reduction in COL1A1^+^, RUNX1^+^, periostin (POSTN)^+^ and Tenascin-C (TNC)^+^ myofibroblasts relative to FT controls (**Fig. 2F-H** and **Supplementary Fig. 2H**).

To assess the impact of disrupting IL-1 signalling in *Fap*-expressing CAFs on PDAC stromal composition, we then analysed PDAC tumours from FIC and FI mice by flow cytometry, Masson’s trichrome staining and multiplex IF analyses. Flow cytometric analysis revealed no significant differences in overall fibroblast abundance, or in immune, endothelial and epithelial cell populations in FIC tumours compared with FI controls (**Supplementary Fig. 2I-L**). However, analysis of CAF subsets showed a significant reduction in iCAF abundance in FIC tumours relative to FI controls, with no significant changes in myCAF or apCAF abundance and no significant difference in the myCAF/iCAF ratio across cohorts (**Fig. 2I**). Consistent with reduced iCAF abundance, PDGFRα⁺αSMA⁻ inflammatory fibroblasts were decreased in FIC tumours relative to FI controls, while collagen deposition and myofibroblast markers were comparable between groups (**Fig. 2J** and **Supplementary Fig. 2M-N**).

Collectively, these results indicate that FTC and FIC models deplete myCAFs and iCAFs, respectively, providing complementary systems to interrogate CAF state-specific functions in PDAC, while revealing broader stromal remodelling following *Tgfbr2* deletion than *Il1r1* deletion in *Fap*-expressing fibroblasts.

### Depletion of myCAFs impairs PDAC liver and lung metastasis formation from primary PDAC tumours

Given the alterations observed in stromal composition in FIC and FTC tumours compared with their respective controls, we next assessed the impact of depleting iCAFs or myCAFs on PDAC progression. Reduced iCAF abundance did not affect primary tumour growth, local dissemination to the diaphragm and peritoneal cavity, or distal dissemination to the liver and lungs relative to FI controls with comparable primary tumour weights (**Fig. 3A-B** and **Supplementary Fig. 3A-B**). Similar to FIC mice, FTC mice showed no difference in primary tumour growth or local dissemination relative to FT controls; however reduced myCAF abundance markedly impaired liver and lung metastasis formation (**Fig. 3C-F** and **Supplementary Fig. 3C-D**).

**Figure 3.**
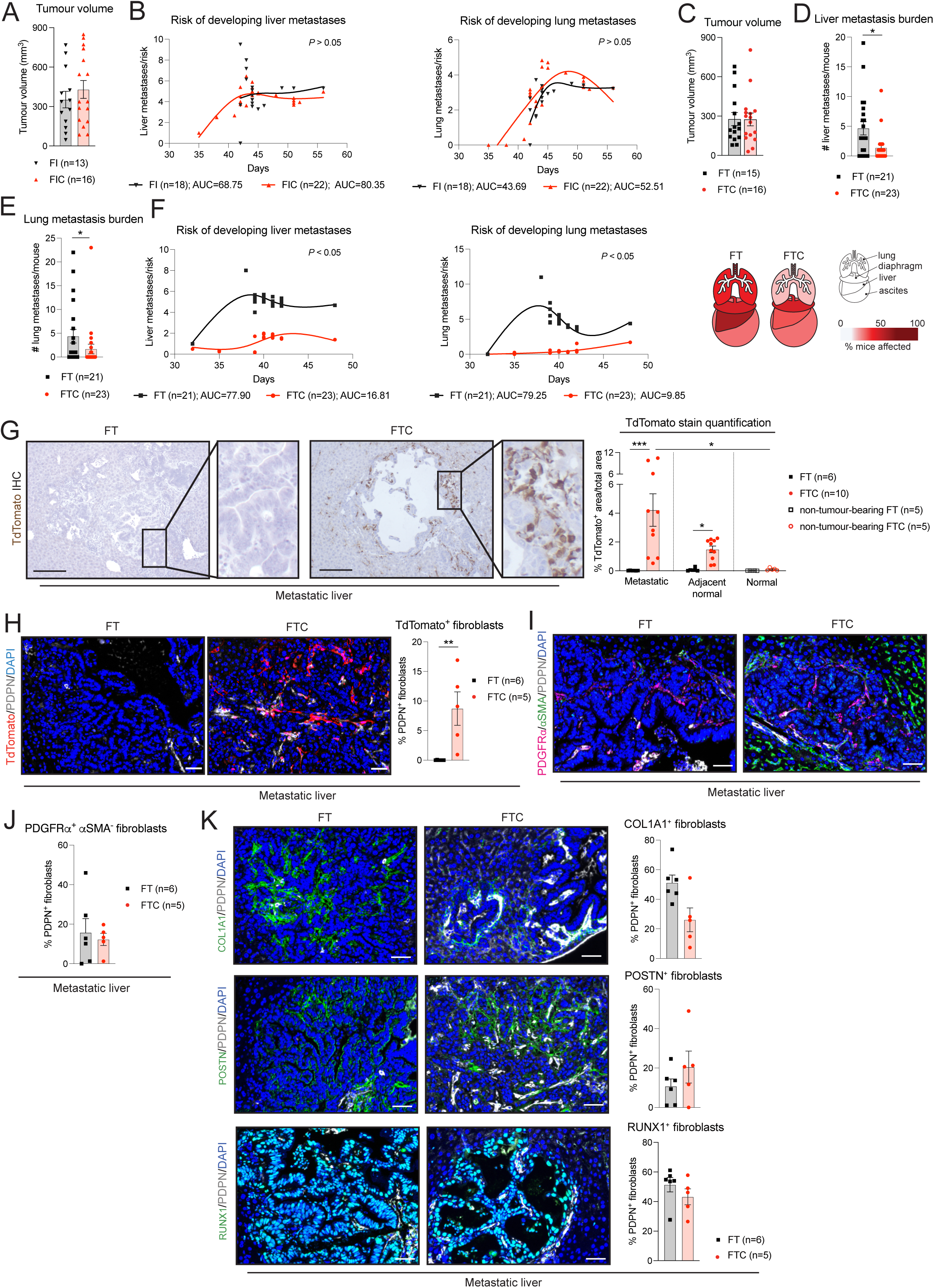
Depletion of myCAFs impairs PDAC liver and lung metastasis formation. All analyses are from OFF dox mice. **(A)** Volumes as measured by ultrasound-based imaging of tumours derived from the transplantation of PDAC organoids in FI (n=13) and FIC (n=16) mice at 36.2 ± 1 (mean ± standard deviation) days post-transplant. Results show mean ± SEM. No statistical difference was found, as calculated by Mann-Whitney test. **(B)** Risk of developing liver and lung metastases in FI (n=18) and FIC (n=22) mice over time as calculated by number of metastases per mouse over time. No significant difference was observed, as calculated by bootstrap test. AUC, area under the curve. **(C)** Volumes as measured by ultrasound-based imaging of tumours derived from the transplantation of PDAC organoids in FT (n=15) and FTC (n=16) mice at 30.5 ± 3.5 (mean ± standard deviation) days post-transplant. Results show mean ± SEM. No statistical difference was found, as calculated by Mann-Whitney test. **(D-E)** Liver **(D)** and lung **(E)** metastasis burden (i.e., number of metastases/mouse) of FT (n=21) and FTC (n=23) mice. * *P* < 0.05, Mann-Whitney test. **(F)** Risk of developing liver and lung metastases in FT (n=21) and FTC (n=23) mice over time as calculated by number of metastases per mouse over time. *P* < 0.05, bootstrap test. Right: schematic summarising the percentages of FT and FTC mice with metastases per anatomical region. **(G)** Left: Representative TdTomato IHC stains of liver metastatic tissues in FT and FTC mice. Scale bars, 100 μm. Inserts, magnifications. Right: TdTomato stain quantification in liver metastatic or adjacent normal tissue from FT (n=6) and FTC (n=10) mice, and in normal liver tissues from non-tumour-bearing FT (n=5) and FTC (n=5) mice. Results show mean ± SEM. * *P adj* < 0.05; *** *P adj* < 0.001; Kruskal-Wallis test. **(H)** Left: Representative mIF images of TdTomato (red), PDPN (white) and DAPI (blue) in FT and FTC liver metastatic tissues. Scale bars, 50 μm. Right: Quantification of TdTomato^+^PDPN^+^CK19^-^DAPI^+^ cells relative to PDPN^+^CK19^-^DAPI^+^ fibroblasts in FT (n=6) and FTC (n=5) liver metastatic tissues. Results show mean ± SEM. ** *P* < 0.01, Mann-Whitney test. **(I)** Representative mIF images of αSMA (green), PDGFRα (pink), PDPN (white) and DAPI (blue) in FT and FTC liver metastatic tissues. Scale bars, 50 μm. **(J)** Quantification of PDGFRα^+^αSMA^-^PDPN^+^CK19^-^DAPI^+^ relative to PDPN^+^CK19^-^DAPI^+^ fibroblasts in FT (n=6) and FTC (n=5) liver metastatic tissues. Results show mean ± SEM. No statistical difference was found, as calculated by Mann-Whitney test. **(K)** Left: Representative mIF images of COL1A1/POSTN/RUNX1 (green), PDPN (white) and DAPI (blue) in FT and FTC liver metastatic tissues. Scale bars, 50 μm. Right: Quantification of COL1A1^+^ or POSTN^+^ or RUNX1^+^ DAPI^+^PDPN^+^CK19^-^ fibroblasts relative to DAPI^+^PDPN^+^CK19^-^ fibroblasts in FT (n=6) and FTC (n=5) liver metastatic tissues. Results show mean ± SEM. No statistical difference was found, as calculated by Mann-Whitney test.

To investigate how myCAF depletion impaired metastasis formation to the liver and lungs, we focused on the FTC and FT models. Because these models disrupt TGF-β signalling in *Fap*-expressing fibroblasts systemically, we first asked whether this targeted stromal lineage was present at liver and lung metastatic sites. TdTomato^+^ cells were detected in both liver and lung metastases and were more abundant than in adjacent non-metastatic tissue from orthotopic PDAC models, or in normal liver and lung tissues from non-tumour-bearing mice (**Fig. 3G** and **Supplementary Fig. 3E-F**). For subsequent analyses, we prioritised liver metastases because they were more frequent than lung metastases in our orthotopic PDAC models, consistent with human PDAC^1^, and showed significant expansion of TdTomato^+^ cells relative to normal liver tissue (**Fig. 3G**). PDPN^+^ fibroblasts were less abundant in liver metastases than in primary tumours, comprising ∼10% of cells within the metastatic TME; within PDPN^+^ fibroblasts ∼10% were TdTomato^+^ (**Fig. 3H and Supplementary Fig. 3G**). In contrast to the stromal remodelling observed in primary FTC tumours, collagen deposition and the abundance of PDGFRα^+^αSMA^-^, COL1A1^+^, POSTN^+^ and RUNX1^+^ fibroblasts were not significantly altered in FTC liver metastases relative to FT controls (**Fig. 3I-K** and **Supplementary Fig. 3H**). Thus, disruption of TGF-β signalling in *Fap*-expressing fibroblasts elicits site-specific stromal responses in PDAC.

Although FTC mice developed fewer spontaneous distant metastases, FTC liver metastases did not show the stromal remodelling observed in primary tumours relative to FT controls. We therefore used experimental models of liver metastasis colonisation and outgrowth to directly test whether disrupting TGF-β signalling in *Fap*-expressing fibroblasts at secondary sites affects metastatic establishment or expansion^43,44^. Intrasplenic injection of PDAC organoids into FTC and FT mice was used to assess liver metastatic colonisation (**Fig. 4A**). Compared with spontaneous liver metastases arising from orthotopic FT PDAC, liver metastases from intrasplenic FT mice showed more pronounced collagen deposition (**Fig. 4B-C**). In the intrasplenic model, collagen deposition was significantly reduced in FTC liver metastases, but not in the adjacent liver parenchyma, indicating that TGF-β-dependent fibroblasts contribute to ECM deposition within experimentally established metastatic lesions (**Fig. 4C** and **Supplementary Fig. 4A**). Although FTC mice showed a lower risk of developing liver metastases than FT controls in intrasplenic models, endpoint metastatic burden was not significantly reduced (**Fig. 4D-E** and **Supplementary Fig. 4B**). We next assessed metastatic outgrowth by injecting PDAC organoids directly into the liver of FTC and FT mice (**Fig. 4F**). As observed in the intrasplenic model, FT intrahepatic tumours showed more pronounced collagen deposition than spontaneous liver metastases from orthotopic FT PDAC, and collagen deposition was significantly reduced in FTC intrahepatic tumours compared with FT controls (**Fig. 4G** and **Supplementary Fig. 4C**). Despite this reduction in ECM deposition, tumour weight and area were unchanged (**Fig. 4H** and **Supplementary Fig. 4D-E**).

**Figure 4.**
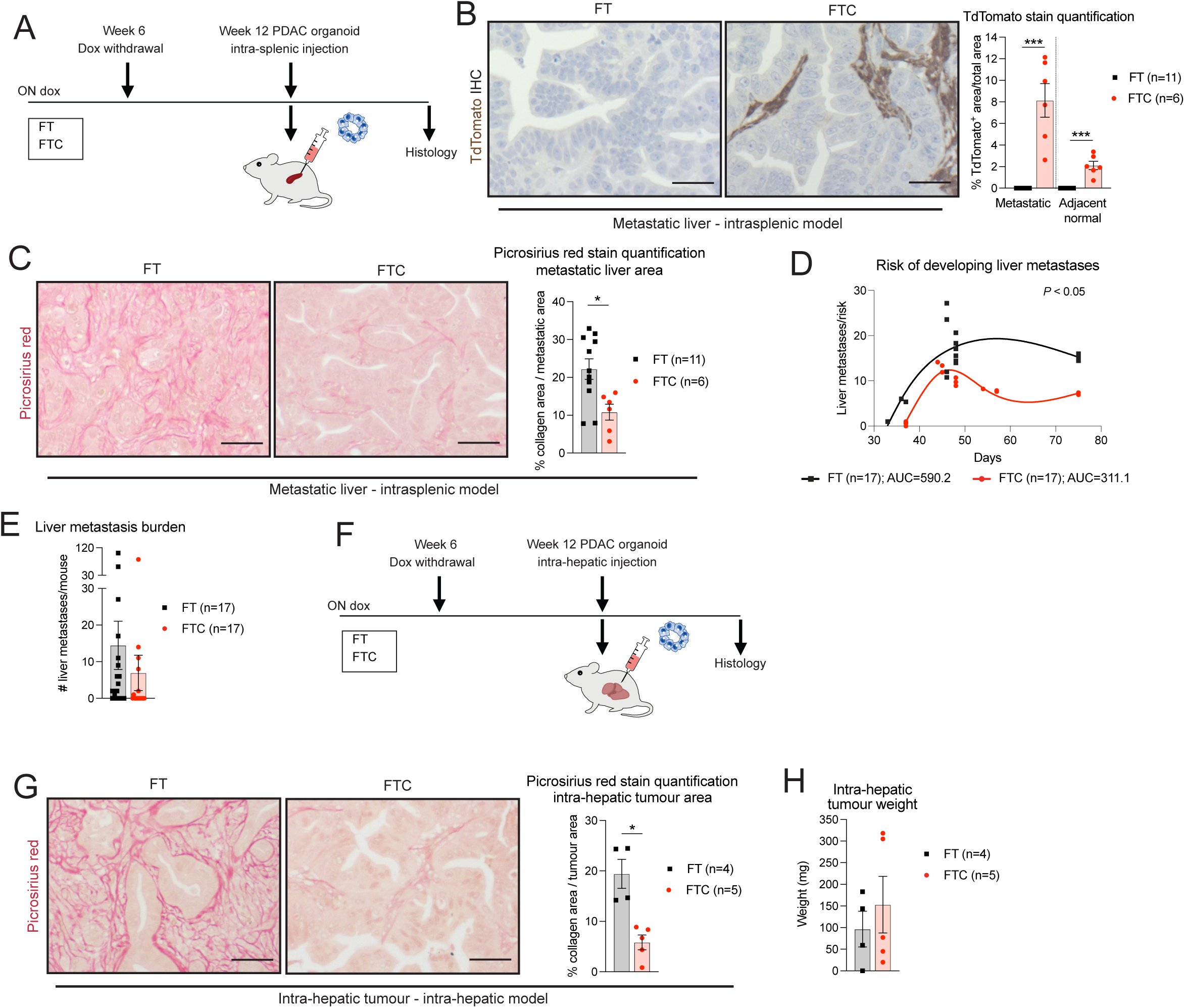
Disruption of TGF-β signalling in *Fap*-expressing fibroblasts does not impair PDAC liver metastatic outgrowth. All analyses are from OFF dox mice. **(A)** Schematic representation of intrasplenic metastasis model. **(B)** Left: Representative TdTomato IHC stains in FT and FTC metastatic liver tissues from intrasplenic models. Scale bars, 25 μm. Right: TdTomato stain quantification in metastatic liver tissues from FT (n=11) and FTC (n=6) intrasplenic models. Results show mean ± SEM. * *P* < 0.05, Kruskal-Wallis test. **(C)** Left: Representative picrosirius red stains of metastatic liver tissues from FT and FTC intrasplenic models. Scale bars, 25 μm. Right: Picrosirius red stain quantification in metastatic liver tissues from FT (n=11) and FTC (n=6) intrasplenic models. Results show mean ± SEM. *** *P adj* < 0.001, Mann-Whitney test. **(D)** Risk of developing liver metastases in FT (n=17) and FTC (n=17) intrasplenic mouse models over time as calculated by number of metastases per mouse over time. *P*, bootstrap test. **(E)** Liver metastasis burden (i.e., number of metastases/mouse) of FT (n=17) and FTC (n=17) intrasplenic mouse models. No statistical difference was found, as calculated by Mann-Whitney test. **(F)** Schematic representation of intra-hepatic metastasis model. **(G)** Left: Representative picrosirius red stains in FT and FTC intra-hepatic tumours. Scale bars, 25 μm. Right: Picrosirius red stain quantification in intra-hepatic tumours from FT (n=4) and FTC (n=5) mice. Results show mean ± SEM. * *P* < 0.05, Mann-Whitney test. **(H)** Weights measured at experimental endpoint of FT (n=4) and FTC (n=5) intra-hepatic tumours at 32.8 ± 0.5 (mean ± standard deviation) days post-transplant. Results show mean ± SEM. No statistical difference was found, as calculated by Mann-Whitney test.

Together, these findings suggest that impaired liver and lung metastasis formation in FTC mice is primarily associated with reduced myCAF-dependent remodelling of malignant cell states within pancreatic tumours, rather than impaired malignant cell outgrowth at secondary sites.

### Depletion of myCAFs drives broad transcriptional changes in the primary TME

To identify primary tumour-associated changes that may underlie the reduced distant metastasis observed in FTC mice, we next performed scRNA-seq of FTC (n=2) and FT (n=2) PDAC tumours (**Fig. 5A** and **Supplementary Fig. 5A-B**; **Supplementary Table 1**). Keeping in line with our multiplex IF findings, CAFs from FTC mice downregulated myofibroblastic markers (e.g. *Postn*, *Runx1*, *Col1a1*) and associated gene signatures (e.g., collagen formation) compared with FT controls, whereas inflammatory programmes (e.g., JAK/STAT signalling) were upregulated (**Fig. 5B-C**; **Supplementary Table 1**).

**Figure 5.**
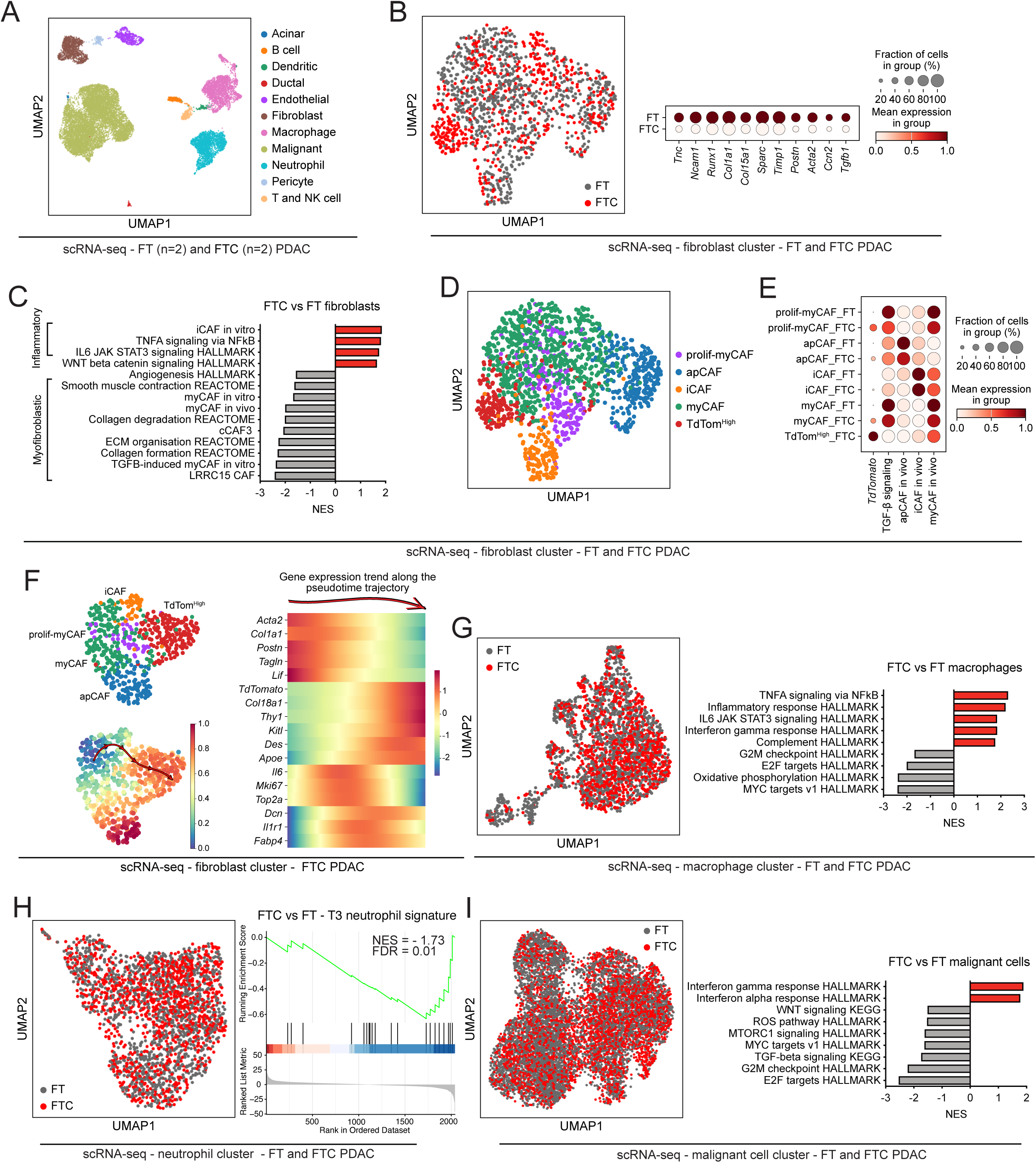
Depletion of myCAFs drives broad transcriptional changes in the primary TME. All analyses are from OFF dox mice. **(A)** Uniform Manifold Approximation and Projection (UMAP) plot of all cells from FT (n=2) or FTC (n=2) PDAC tumours analysed by scRNA-seq. Different cell types are colour coded. **(B)** Left: UMAP of fibroblasts from FT and FTC tumours. Different conditions are colour coded. Right: Dot plot of scaled expression of differentially expressed CAF markers in the fibroblast cluster of FT and FTC tumours. The colour intensity represents the expression level, and the size of the dots represents the percentage of expressing cells. **(C)** Selected significantly upregulated and downregulated pathways identified by gene set enrichment analysis (GSEA) of CAFs from FTC tumours compared to FT tumours, as assessed by MAST analysis from the scRNA-seq dataset. The *in vivo* myCAF signature was obtained from Elyada et al^4^. The *in vitro* iCAF and myCAF signatures were obtained from Öhlund et al^3^. The LRRC15 CAF signature was obtained from Dominguez et al^5^. The cCAF3 signature was obtained from McAndrews et al^15^. NES, normalised enrichment score. **(D)** UMAP of fibroblasts from FT and FTC tumours. Different CAF clusters are colour coded. **(E)** Dot plot of scaled expression of *TdTomato*, the HALLMARK TGF-β signalling pathway and murine i*n vivo* apCAF, iCAF and myCAF signatures in the fibroblast cluster of FT and FTC tumours. The colour intensity represents the expression level, and the size of the dots represents the percentage of expressing cells. The *in vivo* CAF signatures were obtained from Elyada et al^4^. **(F)** Left: UMAP plots of fibroblasts from FTC PDAC showing the CAF clusters (top) and the Palantir-inferred trajectory towards the TdTomato^High^ (TdTom^High^) CAF state in which cells are coloured by Palantir pseudotime and the arrow indicates the inferred direction of state progression (bottom). Right: Heatmap shows the gene expression trends in the TdTom^High^ CAF lineage in the Palantir pseudotime trajectory. **(G)** Left: UMAP of macrophages from FT and FTC tumours. Different conditions are colour coded. Right: Selected significantly upregulated and downregulated pathways identified by GSEA of macrophages from FTC PDAC compared to FT PDAC, as assessed by MAST analysis from the scRNA-seq dataset. **(H)** Left: UMAP of neutrophils from FT and FTC tumours. Different conditions are colour coded. Right: GSEA of T3 neutrophil gene signature in neutrophils from FTC tumours compared to neutrophils from FT tumours, as assessed by MAST analysis from the scRNA-seq dataset. FDR, false discovery rate. **(I)** Left: UMAP of malignant cells from FT and FTC tumours. Different conditions are colour coded. Right: Selected significantly upregulated and downregulated pathways identified by GSEA of malignant cells from FTC PDAC compared to FT PDAC, as assessed by MAST analysis from the scRNA-seq dataset.

To further resolve this complexity, we subclustered fibroblasts and identified previously described iCAFs, myCAFs, apCAFs and proliferating myCAFs (prolif-myCAFs)^42^, together with a distinct TdTomato^High^ CAF population (**Fig. 5D** and **Supplementary Fig. 5C**). Consistent with effective TGF-β pathway disruption, this TdTomato^High^ CAF population showed reduced TGF-β signalling together with lower myCAF-associated signatures and marker expression relative to myCAFs from both FTC and FT tumours (**Fig. 5E** and **Supplementary Fig. 5D**). Notably, myCAFs from FTC tumours also displayed reduced TGF-β signalling and myCAF signatures relative to myCAFs from FT tumours, indicating broader phenotypic remodelling of the CAF compartment beyond the TdTomato^High^ cluster (**Fig. 5E** and **Supplementary Fig. 5D**). Despite these changes, within the TdTomato^High^ CAF cluster, reduced myCAF signature scores were not matched by an equivalent increase in iCAF signature scores (**Supplementary Fig. 5E**). Thus, *Tgfbr2* deletion in *Fap*-expressing fibroblasts does not appear to drive a simple myCAF-to-iCAF transcriptional shift. These data are in agreement with a recent study showing that local deletion of *Tgfbr2* in *Pdgfra*^+^ pancreatic fibroblasts led to a *Col18a1*^High^ CAF state distinct from conventional iCAF and myCAF states^26^. Consistent with this, TdTomato^High^ fibroblasts from FTC tumours were enriched for *Col18a1* expression and the *Col18a1*^High^ CAF gene signature, suggesting that stromal responses to TGF-β signalling disruption converge across distinct fibroblast-targeting strategies (**Supplementary Fig. 5D**and **5F**). TdTomato^High^ CAFs from FTC tumours also upregulated *Thy1*, encoding CD90, but did not downregulate the gene signature associated with the previously described EGFR/ERBB2-activated CD90^-^ metastasis-promoting myCAF subset^8^ (**Supplementary Fig. 5D**and **5F**). This is consistent with the observation that FTC mice showed no change in local dissemination to the diaphragm and peritoneal cavity, the metastatic route primarily linked to CD90^-^ myCAFs^8^. Pseudotime analysis restricted to CAFs from FTC tumours further indicated that TdTomato^High^ CAFs arise from myCAFs, but do not represent a simple intermediate state between myCAFs and iCAFs (**Fig. 5F**). Instead, this trajectory was associated with induction of a distinct transcriptional programme, including upregulation of *Col18a1* and *Thy1* (**Fig. 5F**). Together, these findings underscore the functional complexity of myCAF states and suggest that targeting distinct myCAF populations can have site-specific effects on PDAC dissemination.

Beyond changes in CAF composition, myCAF-depleted PDAC displayed transcriptional alterations in immune and epithelial cell compartments. Macrophages upregulated inflammatory signatures, including TNF signalling and interferon response pathways, while the gene signature of previously described pro-tumorigenic T3 neutrophils^45^ was significantly reduced in FTC tumours relative to FT controls (**Fig. 5G-H**; **Supplementary Table 1**). Malignant cells in FTC PDAC also upregulated interferon response pathways and downregulated TGF-β and WNT signalling, among other signatures, relative to malignant cells in FT PDAC (**Fig. 5I**; **Supplementary Table 1**).

Collectively, these data indicate that depletion of TGF-β-dependent *Fap*-expressing myCAFs induces transcriptional remodelling of stromal, immune and malignant compartments, resulting in a primary TME associated with reduced permissiveness to distant metastatic dissemination. However, because these changes span multiple cellular compartments, directly attributing malignant cell phenotypes to CAF-epithelial crosstalk *in vivo* remains challenging.

### MyCAF-locked PSCs capture TGF-β-dependent CAF signalling observed *in vivo*

Because CAF states are dynamic and can interconvert in response to tumour-derived cues, to resolve CAF-to-malignant cell signalling more directly, we used PSCs, precursors of PDAC CAFs^2,3,10^, engineered to remain ‘locked’ in either a TGF-β-driven myCAF state or an IL-1-driven iCAF state^27^. Briefly, we overexpressed TGF-β1 in CRISPR-engineered *Il1r1* KO PSCs or overexpressed IL-1α in *Tgfbr2* KO PSCs^2,8^ to generate myCAF-locked or iCAF-locked lines, respectively. This strategy constrains CAF plasticity, thus enabling direct dissection of CAF state-specific effects on co-cultured PDAC malignant cells^27^.

We next cultured PDAC organoids alone (monoculture), with both CAF-locked PSC lines (triple co-cultures), or with iCAF-locked or myCAF-locked PSCs (iCAF and myCAF double co-cultures; **Fig. 6A**). Triple co-cultures were included to better recapitulate the *in vivo* setting in which both CAF states co-exist, while comparison with monocultures enabled to distinguish CAF-dependent effects shared by both locked lines from CAF state-specific effects on malignant cell biology. For downstream analyses, co-cultured PDAC cells and CAFs were isolated using a previously established flow-sorting strategy, which separated EpCAM^+^PDPN^-^ PDAC cells from PDPN^+^EpCAM^-^ CAFs^46^ (**Fig. 6A**). To further resolve iCAFs and myCAFs within triple co-cultures, we leveraged our prior RNA-seq data from CAF-locked PSCs cultured in 2D monocultures^27^ and identified *Vcam1* and *Itgb3*, encoding CD106 and CD61, respectively, as differentially expressed surface markers that distinguish PSC-derived CD106^+^CD61^-^ iCAFs from CD61^+^CD106^-^ myCAFs (**Fig. 6A** and **Supplementary Fig. 6A-B**). Consistent with the higher proliferative capacity of myCAFs relative to iCAFs observed *in vivo*^2^, myCAF-locked PSCs expanded approximately two-fold more than iCAF-locked PSCs when co-cultured with PDAC organoids (**Fig. 6B**). Thus, to control for this difference, PDAC organoids were co-cultured with equivalent starting densities of iCAF-locked and myCAF-locked PSCs, or with myCAF-locked PSCs plated at half density. RNA-seq was then performed on flow-sorted PDAC cells from monocultures (n=7), iCAF co-cultures (n=5), myCAF co-cultures (n=9), and triple co-cultures (n=8), as well as on iCAF-locked and myCAF-locked PSCs isolated from double and triple co-cultures. Transcriptomic analysis confirmed that myCAF-locked PSCs retained enrichment of canonical myCAF-associated markers (e.g., *Tgfb1*, *Ctgf*) and signatures (e.g., smooth muscle contraction)^2,3^, and increased activity of SMAD transcription factors (TFs; e.g., *Smad3*) when co-cultured with PDAC organoids alone, irrespective of initial plating density, and when in triple co-cultures with iCAF-locked lines (**Fig. 6C-E** and **Supplementary Fig. 6C-E**; **Supplementary Table 2**). Conversely, iCAF-locked PSCs were enriched for canonical iCAF-associated markers (e.g., *Il1a*, *Cxcl1*) and signatures (e.g., NFκB signalling)^2,3^, and showed increased activity of related TFs (e.g., *Rela*, *Stat3*) across co-culture conditions (**Fig. 6C-E** and **Supplementary Fig. 6C-E**; **Supplementary Table 2**). Together these data confirm stable maintenance of CAF state identity across conditions, enabling direct interrogation of CAF state-specific effects on malignant cells.

**Figure 6.**
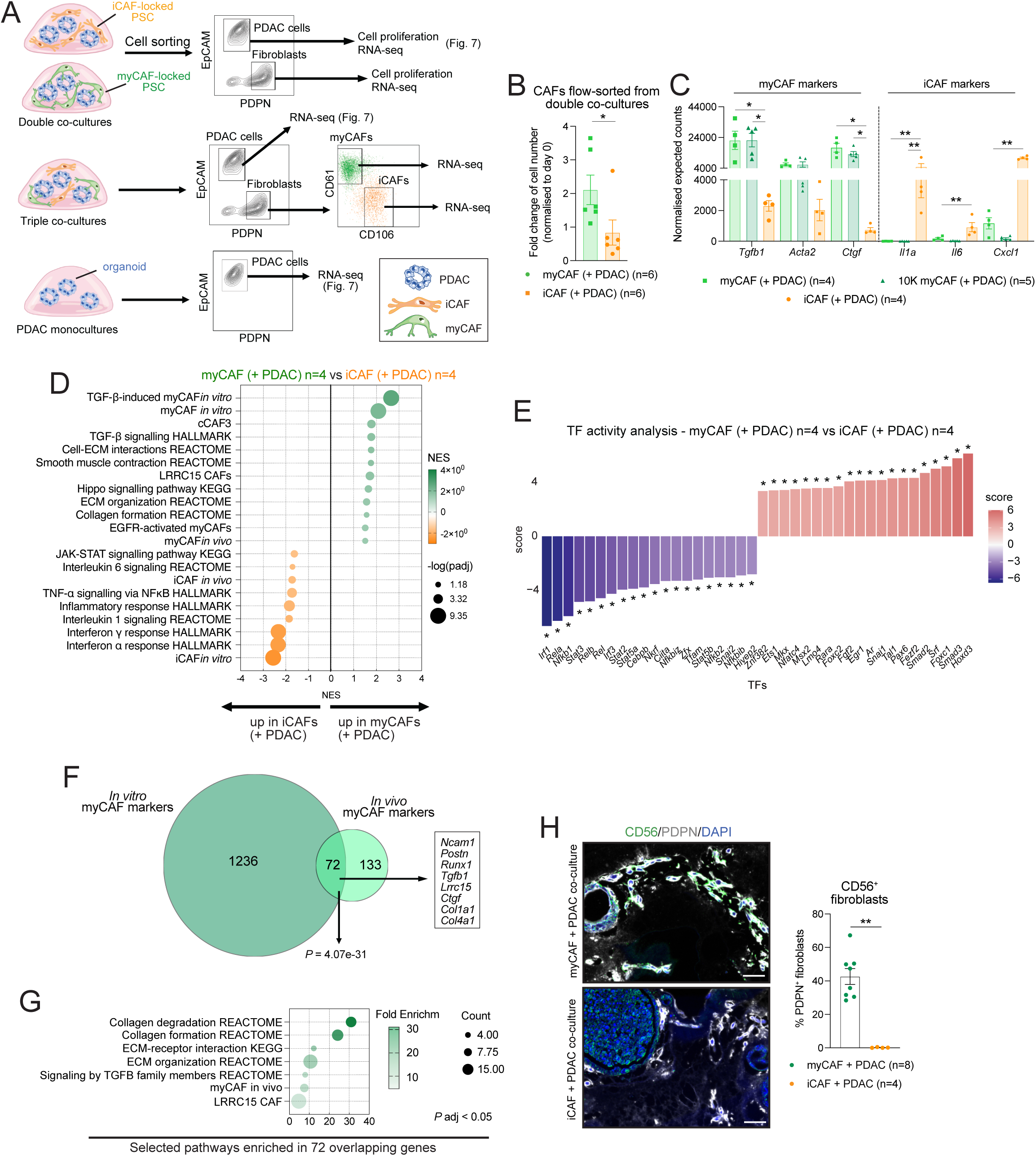
myCAF-locked PSCs capture TGF-β-dependent CAF signalling observed *in vivo*. **(A)** Schematic of flow-sorting strategy of myCAF-locked or iCAF-locked pancreatic stellate cells (PSCs, hereafter myCAFs or iCAFs, respectively) and PDAC organoids from monocultures or co-cultures for RNA-seq analysis. **(B)** Fold change of number of myCAFs (n=6) or iCAFs (n=6) from PDAC organoid co-cultures normalized to day 0. * *P* < 0.05, Mann-Whitney test. For both CAF-locked lines, initial plating was 20K/Matrigel dome. **(C)** RNA-seq expression of myCAF and iCAF markers in myCAFs (n=5 10K myCAFs and n=4 20K myCAFs) and iCAFs (n=4) co-cultured with PDAC organoids. Results show mean ± SEM. * *P adj* < 0.05, ** *P adj* < 0.01, Kruskal-Wallis test. **(D)** Selected significantly upregulated and downregulated pathways identified by GSEA of myCAFs co-cultured with PDAC organoids (n=4) compared to iCAFs co-cultured with PDAC organoids (n=4). The *in vitro* iCAF and myCAF signatures are from Öhlund et al^3^. The TGF-β-induced myCAF *in vitro* signature is from Mucciolo and Araos Henríquez et al^8^. The LRRC15^+^ CAF signature was obtained from Dominguez et al^86^. The *in vivo* iCAF and myCAF signatures are from Elyada et al^42^. **(E)** Activity score of the top 20 most activated and most deactivated transcription factors (TFs) in myCAFs co-cultured with PDAC organoids (n=4) compared to iCAFs co-cultured with PDAC organoids (n=4). * *P* < 0.05, univariate linear model (ulm) method. **(F)** Venn diagrams of genes significantly upregulated (P adj < 0.05) in myCAFs compared to iCAFs co-cultured with PDAC organoids (‘*in vitro* myCAF markers’), and genes significantly downregulated in fibroblasts from FTC tumours compared to fibroblasts from FT tumours (‘*in vivo* myCAF markers’), as assessed by RNA-seq and scRNA-seq, respectively. Significance of the overlap between datasets was defined by hypergeometric test (with denominator = 18,000). **(G)** Significantly enriched pathways (*P* adj < 0.05) based on the 72 genes that are shared by *in vitro* myCAF markers and *in vivo* myCAF markers. **(H)** Left: Representative mIF images of PDPN (white), CD56 (green) and DAPI (blue) stains of PDAC organoid/myCAF co-cultures or PDAC organoid/iCAF co-cultures after 4 days in culture. Scale bars = 50 μm. Right: quantification of CD56^+^PDPN^+^CK19^-^DAPI^+^ cells over PDPN^+^CK19^-^DAPI^+^ cells in PDAC organoid/myCAF co-cultures (n=8) or PDAC organoid/iCAF co-cultures (n=4). Results show mean ± SEM. ** *P* < 0.01, Mann-Whitney test.

We next asked whether this co-culture system could identify TGF-β-dependent myCAF features observed *in vivo*. To this end, we compared bulk RNA-seq datasets from myCAF and iCAF double co-cultures with scRNA-seq datasets from FTC and FT tumours. We identified 1,308 genes with significantly higher expression (*P* adj < 0.05) in myCAFs than in iCAFs when co-cultured with PDAC organoids (hereafter, ‘*in vitro* myCAF markers’) and 205 genes with significantly lower expression in CAFs from FTC tumours relative to FT controls, representing myCAF-associated features lost upon TGF-β pathway disruption *in vivo* (hereafter, ‘*in vivo* myCAF markers’). Of these *in vivo* myCAF markers, 35% (n=72/205) overlapped with *in vitro* myCAF markers (*P*=4.07e-31; **Fig. 6F**; **Supplementary Table 2**). Notably, these 72 shared genes were enriched for TGF-β signalling- and ECM-related pathways and included myofibroblastic markers such as *Ncam1* (encoding CD56)^46^, which we validated at the protein level as significantly upregulated in myCAFs relative to iCAFs when co-cultured with PDAC organoids (**Fig. 6F-H**; **Supplementary Table 2**).

Together, these results support the use of myCAF-locked lines to investigate how TGF-β-dependent myCAF signalling shapes PDAC malignant cell behaviour, providing an *in vitro* platform to identify stromal mechanisms that may underlie the pro-metastatic phenotype observed *in vivo*.

### MyCAF-locked PSCs upregulate EMT signalling in PDAC malignant cells

To define the impact of CAF state-specific signalling on PDAC malignant cells, we analysed PDAC organoids flow-sorted from monocultures, myCAF co-cultures, iCAF co-cultures and triple co-cultures (**Fig. 6A**). PDAC organoids co-cultured with iCAF-locked PSCs upregulated interferon response pathways relative to both PDAC organoids in monocultures and myCAF co-cultures, consistent with prior findings^27^ (**Fig. 7A** and **Supplementary Fig. 7A**; **Supplementary Table 3**). They also upregulated cell cycle programmes and showed increased E2F TF activity, in line with their increased proliferation relative to PDAC organoids from myCAF co-cultures (**Fig. 7A-B** and **Supplementary Fig. 7A-C**; **Supplementary Table 3**). In contrast, PDAC organoids co-cultured with myCAF-locked PSCs were enriched for transcriptional programmes, markers and TF activities associated with metastatic competence, including hepatocyte growth factor receptor (MET), epithelial-to-mesenchymal transition (EMT), MAPK, WNT, and TGF-β signalling, relative to both PDAC organoids in monocultures and iCAF co-cultures (**Fig. 7A** and **Supplementary Fig. 7B-D**; **Supplementary Table 3**)^8,47–50^. These programmes remained elevated in triple co-cultures compared with iCAF co-cultures, indicating that myCAF-derived signals can shape malignant cell transcriptional states even in the presence of iCAFs (**Supplementary Fig. 7E**; **Supplementary Table 3**). Consistent with these transcriptional changes, multiplex IF of PDAC organoid/CAF-locked co-cultures and western blot analysis of PDAC organoids cultured with myCAF- or iCAF-conditioned media (CM) confirmed myCAF-associated upregulation of the EMT marker fibronectin (FN1; **Fig. 7C-D**). Western blot analysis further showed increased activation of TGF-β, EGFR and MAPK signalling in PDAC organoids exposed to myCAF-CM, as indicated by elevated phosphorylated SMAD2/SMAD3 (p-SMAD2/SMAD3), p-EGFR and p-p42/44, respectively (**Fig. 7C**).

**Figure 7.**
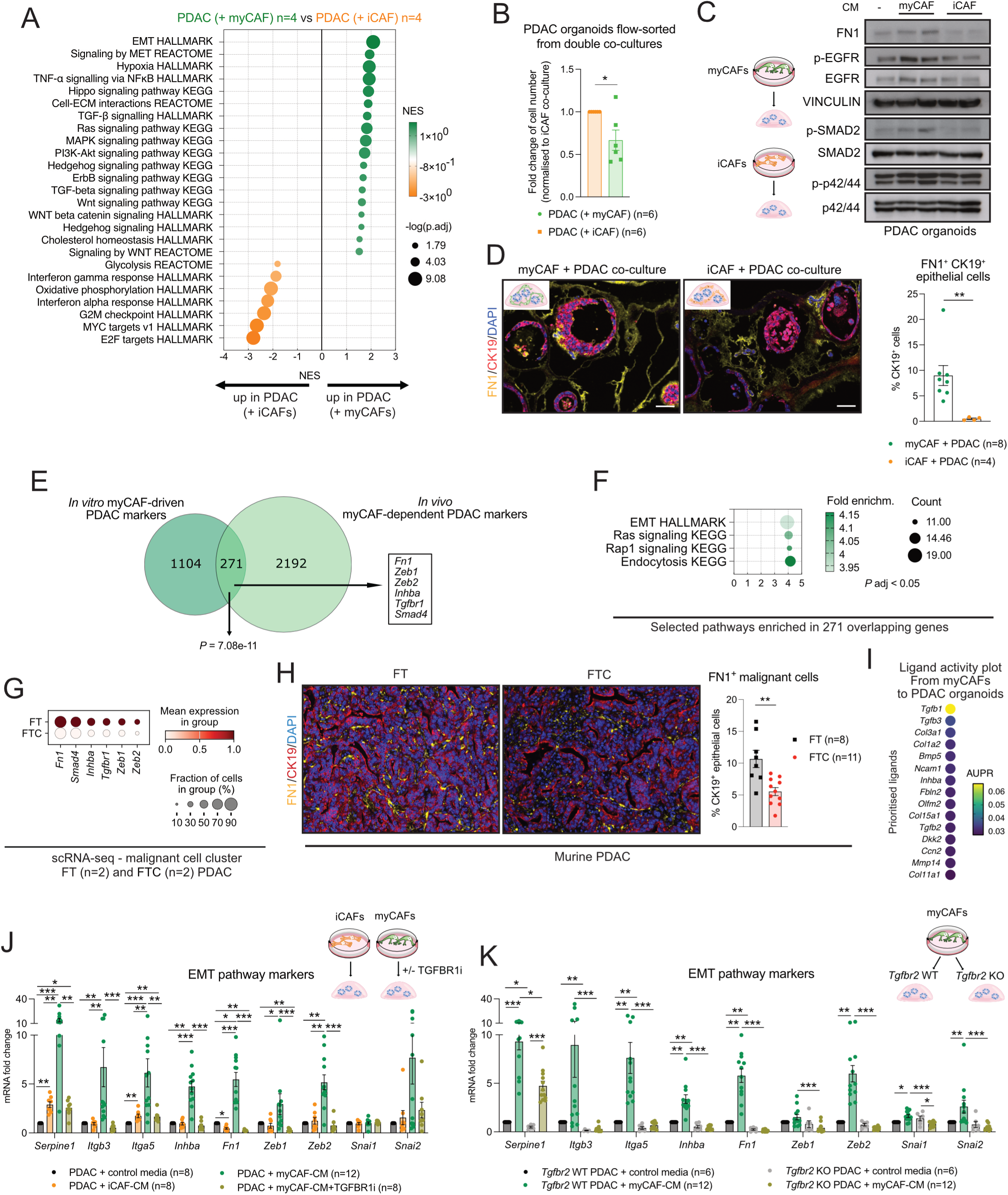
myCAF-locked PSCs upregulate EMT signalling in PDAC malignant cells. **(A)** Selected significantly upregulated and downregulated pathways identified by GSEA of PDAC organoids co-cultured with myCAFs (n=4) compared to PDAC organoids co-cultured with iCAFs (n=4). **(B)** Fold change of number of PDAC organoids from co-cultures with myCAFs (n=6) or iCAFs (n=6) normalized to iCAF co-cultures. * *P* < 0.05, Mann-Whitney test. For both CAF-locked lines, initial plating was 20K/Matrigel dome. **(C)** Western blot analysis of fibronectin (FN1), EGFR, phospho-EGFR (p-EGFR), SMAD2, p-SMAD2/SMAD3, p42/44, p-p42/44 in PDAC organoids cultured with control media (-), with myCAF-conditioned media (CM, n=2) or with iCAF-CM (n=2) for 3 days. VINCULIN, loading control. **(D)** Left: representative mIF images of CK19 (red), FN1 (yellow), and DAPI (blue) stains of PDAC organoid/myCAF co-cultures or PDAC organoid/iCAF co-cultures after 4 days in culture. Scale bars = 50 μm. Right: quantification of FN1^+^CK19^+^PDPN^-^DAPI^+^ over CK19^+^PDPN^-^DAPI^+^ cells in PDAC organoid/myCAF co-cultures (n=8) or PDAC organoid/iCAF co-cultures (n=4), as assessed by mIF. Results show mean ± SEM. ** *P* < 0.01, Mann-Whitney test. **(E)** Venn diagrams of genes significantly upregulated (P adj < 0.05) in PDAC organoids co-cultured with myCAFs compared to PDAC organoids co-cultured with iCAFs (‘*in vitro* myCAF-driven PDAC markers’), and genes significantly downregulated in malignant cells from FTC tumours compared to malignant cells from FT tumours (‘*in vivo* myCAF-dependent PDAC markers’), as assessed by RNA-seq and scRNA-seq, respectively. Significance of the overlap between datasets was defined by hypergeometric test (with denominator = 18,000). **(F)** Significantly enriched pathways (*P* adj < 0.05) based on the 271 genes that are shared by *in vitro* myCAF-driven PDAC markers and *in vivo* myCAF-dependent PDAC markers. **(G)** Dot plot of scaled expression of differentially expressed epithelial-to-mesenchymal transition (EMT) and TGF-β signalling markers in the malignant cell cluster of FT (n=2) and FTC (n=2) PDAC, as assessed by scRNA-seq. The colour intensity represents the expression level, and the size of the dots represents the percentage of expressing cells. **(H)** Left: Representative mIF images of FN1 (yellow), CK19 (red) and DAPI (blue) in FT and FTC PDAC tissues after dox withdrawal. Scale bars, 50 μm. Right: Quantification of FN1^+^CK19^+^PDPN^-^DAPI^+^ epithelial cells relative to CK19^+^PDPN^-^DAPI^+^ epithelial cells in FT (n=8) and FTC (n=11) PDAC tissues after dox withdrawal. Results show mean ± SEM. ** *P* < 0.01, Mann-Whitney test. **(I)** Ligand activity plot of the top ligands in myCAFs inferred to regulate target genes in co-cultured PDAC organoids, as assessed by NicheNet analysis of RNA-seq. AUPR, area under the precision-recall curve. **(J)** qPCR analysis of EMT genes in PDAC organoids cultured with control media (n=8), iCAF-CM (n=8), myCAF-CM (n=12) or myCAF-CM with 1 μM TGF-β receptor 1 inhibitor (TGFBR1i) A83-01 (n=8). * *P adj* < 0.05, ** *P adj* < 0.01, *** *P adj* < 0.001, Kruskal-Wallis test. **(K)** qPCR analysis of EMT genes in *Tgfbr2* WT PDAC organoids cultured with control media (n=6) or myCAF-CM (n=12) and in *Tgfbr2* KO PDAC organoids cultured with control media (n=6) or myCAF-CM (n=12). * *P adj* < 0.05, ** *P adj* < 0.01, *** *P adj* < 0.001, Kruskal-Wallis test.

To narrow down myCAF-driven features of PDAC malignant cells, we applied our cross-model strategy by comparing myCAF-induced transcriptional changes *in vitro* with malignant cell changes associated with myCAF depletion *in vivo*. We identified 1,375 genes with significantly higher expression (*P* adj < 0.05) in PDAC organoids co-cultured with myCAFs than in those co-cultured with iCAFs (hereafter, ‘*in vitro* myCAF-driven PDAC markers’) and 2,463 genes with significantly lower expression in malignant cells from FTC tumours relative to FT controls, representing malignant cell features lost upon myCAF depletion *in vivo* (hereafter, ‘*in vivo* myCAF-dependent PDAC markers’). We found 271 shared genes between these two marker sets, representing a significant overlap (*P*=7.08e-11; **Fig. 7E**; **Supplementary Table 3**). Notably, these shared genes were enriched for EMT and RAS signalling pathways and included EMT-associated markers such as *Fn1*, which we validated at the protein level as significantly downregulated in PDAC malignant cells from FTC tumours relative to FT controls (**Fig. 7F-H**; **Supplementary Table 3**).

To identify candidate myCAF-derived mediators driving the EMT-associated transcriptional changes observed in PDAC organoids, we performed NicheNet analysis on organoid/CAF co-cultures. This analysis inferred TGF-β pathway ligands, including *Tgfb1* and *Tgfb3*, as candidate mediators of myCAF-to-malignant cell communication (**Fig. 7I**). Consistent with this prediction, PDAC organoids cultured with myCAF-CM upregulated EMT-associated genes (e.g., *Itgb3*, *Inhba*, *Fn1*, *Zeb2*) more robustly than those exposed to iCAF-CM, while pharmacological inhibition of TGF-β receptor 1 (TGFBR1) significantly attenuated this response, indicating that myCAF-CM-induced EMT gene expression requires TGF-β pathway activation in malignant cells (**Fig. 7J**). To validate these findings, we generated isogenic *Tgfbr2* wild-type (WT) and *Tgfbr2* knockout (KO) PDAC organoids using CRISPR/Cas9 and co-cultured them with myCAF-CM or control media (**Fig. 7K** and **Supplementary Fig. 7F-G**). Loss of epithelial *Tgfbr2* impaired the induction of EMT-associated genes by myCAF-CM, confirming that malignant cell TGF-β signalling is required for myCAF-mediated EMT induction (**Fig. 7K**).

Together, these data identify myCAF-derived TGF-β signalling as a driver of EMT-associated transcriptional changes in PDAC malignant cells, providing a potential mechanistic link between myCAF state, epithelial plasticity and metastatic potential in primary tumours.

## DISCUSSION

By genetically manipulating two abundant CAF states arising from a shared cell lineage, our study identifies distinct roles for *Fap*-expressing TGF-β- and IL-1-dependent CAFs in shaping primary and metastatic PDAC biology. Depletion of TGF-β-dependent myCAFs remodelled the primary stroma and impaired PDAC metastatic potential to the liver and lungs, with no apparent impact on local metastatic potential to the diaphragm and peritoneal cavity. This phenotype is consistent with previous evidence that PDAC metastatic initiation and progression are shaped by anatomical site-specific pressures^47^. Depletion of myCAFs also appeared to have no impact on primary tumour growth and malignant cell outgrowth in the liver. Together, these data suggest that therapeutic strategies aimed at disrupting TGF-β-dependent myCAF signalling may be most effective at limiting distant metastatic dissemination from primary PDAC, rather than suppressing the outgrowth of already established secondary lesions.

Consistent with the absence of an effect on local metastasis, *Tgfbr2* deletion in *Fap*-expressing CAFs was associated with high *Thy1* expression, encoding CD90, but did not reduce the gene signature defining the EGFR-activated, pro-metastatic CD90^-^ myCAF subset we previously described^8^. Together with the reduced collagen deposition observed in FTC tumours, but not after targeting the CD90^-^ myCAF subset^8^, these findings suggest that perturbing distinct myCAF populations can produce different outcomes in PDAC. Similarly, the reduction in collagen deposition without altered primary tumour growth in FTC mice appears to contrast with prior studies in which depletion of collagen-producing αSMA^+^ myofibroblasts or collagen I accelerated PDAC progression^24,25^. This apparent discrepancy likely reflects differences in the stromal compartments targeted, epithelial-genotype context and disease models and, together with other studies^14,51^, supports the view that ECM-rich, TGF-β-dependent myofibroblastic programmes can exert context-dependent, rather than uniformly tumour-restraining or tumour-promoting, effects. Finally, because TdTomato positivity was also detected in a subset of pericytes and endothelial cells, we cannot exclude the possibility that some changes observed in FTC tumours relative to FT controls involve these cell types. Nevertheless, because our CAF-locked lines are derived from PSCs, our *in vitro* strategy provided a fibroblast-focused approach to isolate TGF-β-dependent myCAF effects on malignant cells, independently of the broader stromal changes observed *in vivo*.

Together with recent findings on *Tgfbr2* deletion in *Pdgfra*^+^ pancreatic fibroblasts^26^, our study indicates that *Tgfbr2* deletion in *Fap*-expressing CAFs leads to stromal transcriptomic reprogramming beyond a simple myCAF-to-iCAF shift. Although *Tgfbr2* deletion in *Pdgfra*^+^ and *Fap*^+^ fibroblasts led to convergent CAF reprogramming, these targeting strategies likely capture only partially overlapping fibroblast populations. Because CAF markers are not universally conserved across organs or disease states^6,19^, complementary fibroblast-specific GEMMs will be needed to disentangle lineage-specific from site-specific CAF functions in primary and metastatic PDAC. Importantly, our off-dox system minimises potential confounding effects of dox exposure, which can influence tumour progression and stromal composition, while avoiding the use of tamoxifen-based induction systems that may also alter the TME^52–54^. It remains to be determined whether *Il1r1* deletion in *Fap*-expressing CAFs similarly induces broader transcriptional remodelling of stromal and immune cell populations beyond the phenotypic changes detected by flow cytometry. More iCAF-rich models, such as *Smad4*-deleted PDAC models^46^, may be particularly useful for dissecting IL-1-dependent CAF functions and revealing context-dependent consequences of iCAF perturbation.

Our understanding of CAF phenotypic and functional heterogeneity at primary tumour sites has expanded substantially in recent years, yet CAF heterogeneity at metastatic sites remains comparatively less well defined. Emerging studies indicate that the same pathway can mark distinct fibroblast states across anatomical contexts: JAK/STAT signalling supports iCAF identity in primary PDAC, but drives myofibroblast activation in liver metastases^2,55^. Extending this concept, our study provides functional evidence that CAF signalling programmes are interpreted in a site-dependent manner: *Tgfbr2* deletion in fibroblasts remodelled CAF composition differently in matched primary tumours and metastases. These findings also highlight the complementary strengths of spontaneous and experimental metastasis models. Orthotopic models better capture the clonal and temporal heterogeneity of liver metastasis formation, as well as systemic conditioning of the liver by a primary pancreatic tumour, including potential pre-metastatic niche formation^56–59^. By contrast, experimental metastasis models provide more controlled settings to assess stromal responses during liver colonisation and outgrowth. For example, the reduction in collagen deposition in FTC lesions in the experimental models, but not in spontaneous orthotopic metastases, suggests that TGF-β-dependent fibroblasts can regulate metastatic ECM deposition in the liver, although this effect may be less apparent in heterogeneous spontaneous metastases. Further analysis of these experimental models may therefore help resolve how TGF-β signalling loss shapes liver metastatic stromal composition.

Our study establishes new GEMMs for CAF state-specific modulation across organs, providing tools to examine how TGF-β- and IL-1-dependent fibroblast programmes operate across disease contexts. Although we focus here on primary tumours and liver metastases, extending these analyses to lungs and other metastatic sites could help distinguish conserved from organ-specific stromal functions. The use of these GEMMs may also extend beyond PDAC, as ECM-producing and inflammatory fibroblasts are also present in pancreatitis^32^, and IL-1 and TGF-β signalling have been implicated in driving iCAF and myCAF formation in other cancers^19,60^. In parallel, the CAF-locked lines and triple co-cultures developed here provide systems to dissect additional CAF state interactions, including iCAF-to-malignant cell crosstalk, reciprocal myCAF-iCAF reprogramming beyond TGF-β and IL-1 signalling, and the combined effects of co-existing CAF states on malignant cell behaviour.

Together with prior studies linking EMT to PDAC dissemination and showing that TGF-β and CAF-derived cues regulate epithelial plasticity^8,28,61–64^, our work identifies myCAFs as candidate stromal mediators of EMT-associated programmes in PDAC malignant cells. While EMT was the focus of our mechanistic studies, the models developed here also provide platforms to define additional CAF state-dependent pathways that may contribute to PDAC progression, including myCAF-induced MAPK and MET signalling^49,65^. More broadly, our study supports a framework for stage- and site-informed stromal targeting in PDAC, in which CAF state, anatomical context and timing of intervention are considered together.

## MATERIAL AND METHODS

### Mouse models

Males and females C57BL/6J mice were purchased from the Charles River Laboratory (strain number 632). C57BL/6J-background *Tgfbr2^fl/fl^* mice (strain #012603, RRID:IMSR_JAX:012603)^33^ and *Il1r1*^fl/fl^ mice (strain #028398, RRID: IMSR_JAX:028398)^40^ were purchased from JAX. C57BL/6J-background *Rosa26^LSL-tdTom/tdTom^* mice were kindly shared by Professor Winton (previously at CRUK Cambridge Institute, CRUK-CI). The C57BL/6J-background Tg (*Fap*-tTA) mouse line was kindly shared by Professor Fearon (previously at CRUK-CI)^30^. The C57BL/6J-background tet-O-Cre-LC1 mouse line (Tg(tetOcre)LC1Bjd; European Mutant Mouse Archive) was kindly shared by Professor Linterman (Babraham Institute and Malaghan Institute of Medical Research)^30,66^. Specific qPCR probes for *Fap*-tTA, tet-O-Cre, *Rosa26*, *Tgfbr2* and *Il1r1* were developed and genotyping assays were run by Transnetyx. All animals were housed in accordance with the guidelines of the UK Home Office “Code of Practice for the Housing and Care of Experimental Animals”. They were kept behind strict barriered housing, fed expanded rodent diet (Labdiet) and filtered water ad libitum. All animal procedures and studies were reviewed by the CRUK-CI Animal Welfare Ethical Review Body (AWERB), approved by the Home Office and conducted under PP4778090 and PP7672420 project licences in accordance with relevant institutional and national guidelines and regulations.

### Pancreatic orthotopic transplantation models

*Fap*-tTA; *Rosa26^LSL-tdTom/tdTom^; Tgfbr2^fl/fl^* controls (i.e., FT mice), *Fap*-tTA; tet-o-Cre; *Rosa26^LSL-tdTom/tdTom^; Tgfbr2^fl/fl^* (i.e., FTC mice), *Fap*-tTA; *Rosa26^LSL-tdTom/tdTom^; Il1r1^fl/fl^* controls (i.e., FI mice) and *Fap*-tTA; tet-o-Cre; *Rosa26^LSL-tdTom/tdTom^; Il1r1^fl/fl^* (i.e., FIC mice) were kept on dox water until ∼6-7 weeks of age (0.2 g/L dox in water, D9891; Sigma-Aldrich). We observed that removing dox before this timepoint was associated with abnormal gait and weight loss in FTC mice. By contrast, FIC mice showed no obvious clinical signs even when dox was removed before 6 weeks of age. Orthotopic injections were conducted as previously described^36^. Briefly, 10,000 single cells prepared from organoid cultures were resuspended as a 35 μL suspension of 50% Matrigel in PBS and injected into the pancreas of males and females ∼13-to-15-week-old (∼3 months-old) mice. Pancreatic tumours were imaged using the Vevo 2100 Ultrasound at two different orientations with respect to the transducer. Tumour volumes were measured at two angles using the Vevo LAB software program (version 5.7.0). Tumour volume analyses were performed blindly prior to plotting the data for visualisation. Only mice with successful (i.e., non-leaked/re-injected) orthotopic injections were included for tumour volume and metastasis injections.

### Intrasplenic models

Intrasplenic injections were conducted as previously described^44^. Briefly, 20,000 single cells prepared from organoid cultures were resuspended as a 50 μL suspension in PBS and injected into the spleen of male and female ∼12-to-16-week-old (∼3 months-old) mice.

### Intrahepatic models

Intrahepatic injections were conducted as previously described^43^. Briefly, 10,000 single cells prepared from organoid cultures were resuspended as a 10 μL suspension of 50% Matrigel in PBS and injected into the liver of female ∼13-to-18-week-old (∼3-4 months-old) mice.

### Quantification of metastases

Metastases were counted from one H&E-stained section for each liver and lung tissue using Aperio ImageScope or QuPath softwares. Analysis of cohorts was performed blindly. The risk of developing metastasis over time was calculated by dividing the cumulative number of metastases for the total number of mice at each time point. Bootstrap test was used to calculate the p value between the two curves.

### PCR of Tgfbr2 and Il1r1

To validate gene knockout in FTC and FIC mouse models, individual PCRs were set up to amplify the floxed exon 4 of *Tgfbr2*, and the floxed exons 3 and 4 of *Il1r1*, respectively. Primers used were published previously^33,40^. *Tgfbr2* PCR was performed on TdTomato^High^ and TdTomato^-^ fibroblasts from off-dox FTC PDACs, as well as on fibroblasts and malignant cells from off-dox FT PDACs and on-dox FTC PDACs. *Tgfbr2* PCR was performed in a 20 μL reaction using 0.5 μM of each primer (*Tgfbr2_In3 F1: TATGGACTGGCTGCTTTTGTATTC; Tgfbr2_In3 R1: TGGGGATAGAGGTAGAAAG ACATA; Tgfbr2_In4 R2: TATTGGGTGTGGTTGTGGACTTTA*) and 10 ng of genomic DNA, using the following PCR conditions: 98°C for 30 s; 35 cycles of 98°C for 10 s, 66°C for 30 s, 72°C for 40 s; 72°C for 2 min; 4°C hold. *Il1r1* PCR was performed on TdTomato^+^ fibroblasts (i.e., also including TdTomato^Low^ cells), TdTomato^-^ fibroblasts and malignant cells from off-dox FIC PDACs, as well as on fibroblasts and malignant cells from off-dox FI PDAC. *Il1r1* PCR was performed in a 20 μL reaction using 0.5 μM of each primer (*Il1r1_In2 F1: CTTGTGTCCTATGGGTGTCC; Il1r1_In4 R1: GTACCAATGGAGGCCAGAAG*) and 20 ng of genomic DNA, using the following PCR conditions: 98°C for 30 s; 35 cycles of 98°C for 10 s, 66°C for 30 s, 72°C for 30 s; 72°C for 2 min; 4°C hold. Q5 High-Fidelity DNA polymerase (M0491S, New England Biolabs) was used for both PCRs. PCR products were separated in a 1.5% agarose gel containing SYBR Safe stain (S33102; Invitrogen) in TBE (Tris-Borate-Ethylenediaminetetraacetic acid) buffer. Gel imaging was performed using a Syngene UV transilluminator. PCR gel bands were quantified using Image J. Briefly, bands of interest were selected using the rectangle tool and assigned to lanes, then a stacked graph of peak plots representing band intensity was generated using the gel analyse tool. The intensity of the band was converted into an area using the line tool to select the band peak and wand tool to calculate the area. The area represents the intensity of the band.

### Rheology

Pancreatic tumour tissues were harvested in ice-cold PBS + 1% BSA. Cylindrical punches of ∼6 mm diameter were cut from the bulk tumour using a biopsy punch (WB100040, Qiagen). The samples were gently blotted on filter paper to remove excess surface buffer, snap frozen in liquid nitrogen and stored at -80°C until rheological analysis. The samples were thawed at room temperature and mounted on the lower 20 mm sandblasted Peltier plate (519080.906, TA Instruments) for rheological analysis using an 8 mm sandblasted upper plate geometry on a HR2 hybrid rheometer (533002.901, TA Instruments). The measurement gap was adjusted to 1000 µm and a small normal force threshold of (0.01–0.1 N) was used to ensure good contact. The excess buffer was blotted from the tissue while loading, and the tissue was trimmed from the plate edges after setting the gap. All measurements were performed with the temperature control set to 37°C, using a Peltier plate and a humidity cover to prevent evaporation. Oscillatory time sweeps with a temperature soak of at least 30 s, an experimental time of 60-120 s, a strain of 0.5%, and an angular frequency of 1 rad/s were run in between experiments.

#### Frequency sweep

Frequency-dependent viscoelastic properties were measured using oscillatory frequency sweeps at a constant strain amplitude of 0.5%, with angular frequency varied from 0.1-100 rad/s. The storage modulus (G′) and loss modulus (G″) were recorded and the complex viscosity (η*) was calculated. Higher storage modulus (G′) values indicate greater elastic stiffness (i.e., greater resistance to mechanical deformation). For visualisation, measurements from individual biological replicates were summarised as mean ± standard error of the mean (SEM) at each angular frequency. For statistical analysis, *G′*, *G″*, and *η\** values at angular frequencies of 0.1, 1, and 10 rad/s were extracted from each individual biological replicate and compared between FT and FTC groups using unpaired two-sided Welch’s t-tests.

#### Amplitude sweep

To determine the linear viscoelastic region (LVR) and ensure the strain amplitude for frequency sweep experiments is within this LVR, an oscillatory amplitude sweep was performed at a fixed angular frequency of 1 rad/s over a strain range of 0.1–500%. The amplitude sweep was run from low to high strain to minimise pre-damage, and both the storage modulus (G′) and loss modulus (G″) were recorded. For visualisation, individual biological replicate curves were summarised as mean ± SEM at each strain value. The LVR was identified as the region in which the storage modulus (G′) showed minimal variation with increasing strain. In FT samples, G′ and G″ began to decrease with increasing strain above 0.5 %, indicating the onset of strain-dependent changes in the viscoelastic response. A strain amplitude of 0.5% was therefore selected for subsequent frequency sweeps as a conservative strain within the LVR, minimising potential strain-induced structural changes during measurement. For statistical analysis, G′ and G″ values at 0.1%, 1%, and 10% strain were extracted from each individual biological replicate and compared between FT and FTC groups using unpaired two-sided Welch’s t-tests.

### Cell lines and cell culture

Murine PDAC KPC organoid lines T69A (female) and T6-LOH (male) were previously described and cultured in organoid complete media unless otherwise specified, as previously described^3,67,68^. Organoids were cultured for no more than 30 passages at 37°C with 5% CO2. For myCAF-locked lines: TGF-β-expressing PSC4 (G1, E8) and PSC5 (F3, C9, C3) *Il1r1* KO lines from Biffi et al 2019^2^. For iCAF-locked lines: IL-1α-expressing PSC4 (A3, A10) and PSC5 (B11, D4, E8) *Tgfbr2* KO from Mucciolo et al 2024^8^. These locked lines cells have also been used in Pelicano et al^27^. To note, these cell lines are hygromycin resistant (pBABE-hygro), puromycin resistant (SV40 plasmid for immortalization) and neomycin resistant (LR*N*G plasmid for *Il1r1* or *Tgfbr2* KO). They are also GFP-positive (LRN*G* plasmid for *Il1r1* or *Tgfbr2* KO). Mycoplasma testing for cell lines was performed prior to each freezing.

### *Tgfbr2* CRISPR/Cas9 knockout in PDAC organoids

To knock out *Tgfbr2* in KPC T6-LOH and T69A organoid lines, a CRISPR guide (phosphorothionate-modified sgRNA, 456: TTCATGCGGCTTCTCACAGA) was designed against exon 2 of the murine *Tgfbr2* gene (ENSMUST00000035014.8). A guide which targets the *Rosa26* locus was also included to generate control (i.e. wild-type) lines (265: GAAGATGGGCGGGAGTCTTC). Mouse PDAC organoids were dissociated into single cells and 100,000 cells were electroporated using an Amaxa 4D Nucleofector unit (Lonza) with 4 μg TrueCut spCas9 protein V2 (A36498; Invitrogen) and 80 pmol guide RNA (Synthego), using program CM-137 and P3 nucleofector solution (V4XP-3032; Lonza). A cell pellet was taken 3- and 10-days post electroporation, and genomic DNA was extracted using the DNeasy blood and tissue kit (69506; Qiagen). Exon 2 of *Tgfbr2* was amplified by PCR using the Q5 High Fidelity DNA polymerase (M0491S; NEB). *Tgfbr2* primers used were FWD: TGCGTGCATCCAGGAATTCT, and REV: TTGATGTCCAGCTGCTCCAG. Amplicons were subjected to Sanger sequencing and analyzed using Synthego ICE web tool to calculate the percent editing in a pool. Organoids were plated as single clones. Eleven clones were pooled together to generate each *Tgfbr2* WT organoid line. Knockout was confirmed by western blot analysis, and clones were pooled together to generate *Tgfbr2* KO lines (6 clones/T6-LOH KPC organoid line; 8 clones/T69a KPC organoid line).

### Western blot

Organoids were harvested in Cell Recovery Solution (354253; Corning) supplemented with complete, mini protease inhibitors (11836170001; Roche) and a phosphatase inhibitor cocktail (4906837001; Roche) and incubated for 30 min at 4°C. Cells were pelleted at 1500 rcf for 5 min and lysed in 0.1% Triton X-100, 15 mmol/L NaCl, 0.5 mmol/L EDTA, 5 mmol/L Tris, pH 7.5, supplemented with complete, mini protease inhibitors (11836170001; Roche) and a phosphatase inhibitor cocktail (4906837001; Roche). Cells were incubated on ice for 30 min briefly vortexed and pelleted at 13,200 rpm for 10 min at 4°C. Concentration of protein collected in the supernatant was determined using DC protein assay (5000113-5; Bio-Rad). Standard procedures were used for western blotting. Primary antibodies used were ACTIN (8456; Cell Signaling Technology; RRID:AB_10998774), FN1 (ab2413; Abcam RRID:AB_2262874), p-EGFR (3777; Cell Signaling Technology; RRID: AB_2096270), EGFR (4267; Cell Signaling Technology; RRID: AB_2246311), VINCULIN (13901: Cell Signaling Technology; RRID:AB_2728768), p-SMAD2/SMAD3 (8828; Cell Signaling Technology; RRID: AB_2631089), SMAD2 (5339; Cell Signaling Technology; RRID: AB_10626777), p-p42/44 (4370; Cell Signaling Technology; RRID: AB_2315112), p42/44 (4695; Cell Signaling Technology; RRID: AB_390779) and TGFBR2 (sc-17792; Santa Cruz Biotechnology; RRID: AB_628349). Proteins were detected using appropriate HRP-conjugated secondary antibodies (Jackson ImmunoResearch Laboratories).

### *In vitro* organoid treatments

PDAC organoids were treated with 2 ng/mL human TGF-β1 (T7039-2UG; Sigma) or iCAF conditioned media (CM), or myCAF CM with or without 1 μM A83-01 (SML0788; Sigma) in reduced media (i.e. 5% FBS DMEM) for 72 hours prior to collecting protein or RNA.

### Reverse transcription quantitative polymerase chain reaction analyses

RNA (up to 1 μg) of PDAC organoids was reverse transcribed using TaqMan reverse transcription reagents (N808 0234; Applied Biosystems). qPCR was performed using gene-specific TaqMan probes (Thermo Fisher Scientific) and TaqMan master mix (4440040; Applied Biosystems) on a QuantStudio 6 Flex Real-Time PCR system. Gene expression was normalized to *Hprt* and calculated using the 2^–ΔΔCt^ method.

### Immunohistochemical and histological analyses

Standard procedures were used for immunohistochemistry (IHC). Primary antibody for IHC was anti-DsRed (for detecting TdTomato; 632496; Takara Bio; RRID:AB_10013483). Hematoxylin (H-3404; Vector Lab) was used as nuclear counterstain. Hematoxylin and eosin and Masson’s trichrome stains were performed according to standard protocols by the Histology core at the CRUK-CI. Standard procedures were used for picrosirius red staining. Brightfield images of tissue slides were obtained with an Axio Vert.A1 (ZEISS). Stained sections were scanned with Aperio ScanScope CS and analysed using the ImageScope Positive Pixel Count algorithm. The percentage of collagen area was then determined by calculating the percentage of blue (for Masson’s trichrome stain) or red (for picrosirius red stain) pixels relative to the entire tissue. To quantify TdTomato stains, the percentage of positive pixels was calculated relative to the entire tissue. Quantifications were performed blindly prior to plotting the data for visualisation.

### Multiplex immunofluorescence analyses

Paraffin embedded tissue sections were dewaxed and rehydrated as per standard procedures on an automated multistainer (Leica ST5020) then washed in deionized water. Tissue sections went through five rounds of antigen retrieval (AR), followed by blocking, incubation with primary antibody then secondary antibody and finally opal fluorophore. Briefly, AR was then performed by immersing the tissue slides in AR pH9 1X Tris-EDTA buffer (Abcam, ab93684) or AR pH6 1X AR6 buffer (Akoya Biosciences, AR600250ML) and boiling in a microwave for 20 min. After cooling, tissue slides were washed in deionized water by rocking twice for 2 min and then a further 2 min washing in PBS + 0.1% Tween20 (PBS-T). Tissue sections were then outlined using a hydrophobic barrier pen and blocking buffer (Akoya Biosciences, ARD1001EA) was incubated on the slides at room temperature (RT) for 15 min in a humidified chamber. Slides were incubated with primary antibody for 45 min at RT or 4°C overnight. Washing was performed by rocking the slides in PBS-T for 3 x 2 min. This was followed by incubation with secondary antibody (Akoya Biosciences, ARH1001EA) for 15 min at room temperature, washing, then incubation with opal fluorophore for 12 min. Slides were washed again then immersed in antigen retrieval buffer to being the cycle from microwaving to fluorophore incubation for all other primary antibodies in the panel. After the last round of staining incubation with opal fluorophore 780, DAPI was added to tissue sections (Akoya Biosciences, FP1490) for 5 min, and to co-culture sections for 30 min (Merk, D8417). After washing, the slides were then mounted in ProLong Gold Antifade mountant (Thermofisher, P36930) and 1.5 mm coverslip (2980-245, Corning). The slides were then imaged using the Vectra Polaris. Tissue staining was analyzed using QuPath software. For each round of staining, all sections were set to the same signal thresholds per channel, annotated by area to analyze and then cells detected using DAPI signal to identify each nuclei using the cell detection tool. Positivity for the markers of interest was defined by signal threshold in the object classifer tool and all parameters applied to each section remained the same. Primary antibodies used to identify these markers are as follows: αSMA (ab5694; Abcam; RRID:AB_2223021), PDPN (127403; BioLegend; RRID: AB_1134221), PDGFR-α (3174; Cell Signaling Technology; RRID: AB_2162345), FN1 (Ab2413; Abcam; RRID:AB_2262874), CD56 (99746; Cell Signaling Technology; RRID:AB_2868490), TdTomato (632496; Takara Bio; RRID:AB_10013483), TNC (ab108930; Abcam; RRID:AB_10865908), RUNX1 (MA5-50434; thermos Fisher Scientific; RRID:AB_3094219), COL1A1 (72026; Cell Signaling Technology; RRID:AB_2904565), CD106 (ab134047; Abcam; RRID:AB_2721053), CD61 (ab179473; Abcam; RRID:AB_2917988), POSTN (ab227049; Abcam; RRID:AB_3095822), ECAD (610182; BD Biosciences; RRID:AB_397581) and CK19 (TROMA-III; DSHB; RRID: AB_2133570). All buffers, anti-rabbit and anti-mouse secondary antibodies, opal dyes and DAPI used were supplied in the Opal 6-Plex manual dection kit (Akoyabiosciences, NEL861001KT) unless otherwise detailed, additional goat-anti rat secondary (Vector, MP-7444-15) was used for antibodies raised in rat, and additional goat anti-Syrian hamster secondary (ab6892; Abcam; RRID:AB_955427) was used for antibodies raised in Syrian hamster.

### Flow cytometry

Tumours were processed as previously described^2^. Cells were blocked for 15 minutes on ice with CD16/CD32 Pure 2.4G2 (553142, BD Bioscience). For flow-cytometric analysis of CAFs, endothelial cells, epithelial cells and immune cells, cells were stained for 30 minutes on ice with anti-mouse CD31-PE/Cy7 (102418; BioLegend; RRID:AB_830757), CD45-PerCP/Cy5.5 (103132; BioLegend; RRID:AB_893344), CD326 (EpCAM)-AlexaFluor 488 (118210; BioLegend; RRID:AB_1134099), PDPN-APC/Cy7 (127418; BioLegend: RRID:AB_2629804), MHCII-BV785 (107645; BioLegend; RRID:AB_2565977), and Ly6C-APC (128015; BioLegend; RRID:AB_1732087).

For flow-cytometric analysis of lymphocytes, cells were stained for 30 minutes on ice with anti-mouse CD19-PE/Cy7 (115520; BioLegend; RRID: AB_313655), CD45-PerCP/Cy5.5 (103132; BioLegend; RRID:AB_893344), TCR-β-Alexa488 (109215; BioLegend; RRID:AB_493344), CD3e-Alexa488 (100321; BioLegend; RRID:AB_389300), CD8-APC/Cy7 (100713; BioLegend; RRID:AB_312752), and CD4-APC (100515; BioLegend; RRID:AB_312719).

For flow-cytometric analysis of macrophages and neutrophils, cells were stained for 30 minutes on ice with anti-mouse CD45-PerCP/Cy5.5 (103132; BioLegend; RRID:AB_893344), CD11b-PE/Cy7 (101215; BioLegend; RRID:AB_312798), F4/80-BV785 (123141; BioLegend; RRID:AB_2563667), MHCII-APC/Cy7 (107627; BioLegend; RRID:AB_1659252 – data not shown) and Gr1-APC (108411; BioLegend; RRID: AB_313376).

Cells were resuspended in PBS with DAPI and analysed on a BD FACSymphony cell analyser. Flow analyses were performed blindly prior to plotting the data for visualization.

### Cell sorting of PDAC organoid/PSC co-cultures for RNA-sequencing

For double co-cultures, organoids were split into two equal parts for co-culture with iCAF-locked or myCAF-locked PSCs, so that organoid proliferation could be compared between the two types of co-cultures plated on the same day. Organoids were embedded in Matrigel with 20,000 (or 10,000 where specified) CAF-locked PSCs per dome. Sorting of PDAC organoid/PSC co-cultures was performed after 3.5 days culture in reduced media (i.e. 5% FBS DMEM). Following single cell digestion of co-cultures, cells were stained for 30 min on ice with anti-mouse CD326 (EpCAM)-PE (118205; BioLegend; RRID:AB_1134176) and PDPN-AlexaFluor 488 (156208; BioLegend; RRID:AB_2814080). Cells were resuspended in PBS with DAPI and sorted with a BD FACSMelody cell sorter. Sorted cell numbers were used for quantifying proliferation during co-culture.

### RNA-sequencing analyses of PDAC organoids and PSCs flow-sorted from co-cultures and monocultures

RNA-seq data of PDAC organoid monocultures and co-cultures are available at the Gene Expression Omnibus (GEO) under the accession number GSE342729. RNA-seq analyses are included in **Supplementary Tables 2-3**. Samples were collected in 1 mL of TRIzol Reagent (15596018; Invitrogen). RNA was extracted using the PureLink RNA mini kit (12183018A; Invitrogen). RNA concentration was measured using a Qubit RNA Broad Range (Q10210; Invitrogen) or High Sensitivity (Q32852; Invitrogen) Assay kit and RNA quality was assessed on a TapeStation 4200 (G2991BA; Agilent Technologies) using the TapeStation High Sensitivity RNA ScreenTape (5067-5579; Agilent Technologies) analysis. mRNA library preparations were performed using 95-300 ng per sample (RNA integrity number, RIN > 7.9). Illumina libraries were then sequenced on NovaSeq6000. Raw FASTQ files were aligned to the GRCm39 (release 109, kmer 23) reference genome using the quasi-alignment tool Salmon^69^. The resulting transcript-level estimates were aggregated to the gene level using the R package tximport^70^. Samples were combined and genes with fewer than 10 total counts across all samples were excluded from downstream analyses. DESeq2 R package was used for differential expression analysis (DEA)^71^. Genes with an adjusted P value < 0.05 were considered significantly differentially expressed. Genes were pre-ranked based on a scoring metric: −log10(P value) × sign(log2(fold change)) and gene Set Enrichment Analysis (GSEA) was performed using the R package clusterprofiler^72^ against the Hallmark, Reactome, and C2 canonical pathway collection (C2.cp.v5.1) downloaded from the Molecular Signatures Database (MSigDB)^73^. Pathways were considered significantly enriched at a false discovery rate (FDR) < 0.25 and an absolute normalized enrichment score (|NES|) > 1.5. NichenetR^74^ was applied to bulk RNA-seq. Genes of interest were defined from significant DEGs (Padj < 0.05, |log2fold change| > 0) and all expressed genes were considered as background genes. Transcription factor activity analysis was performed using a univariate linear model (ULM) as implemented in decouplerR package^75^ with regulons from CollecTRI network^76^. The stat column from the DESeq2 output was used as input. TFs with p-value < 0.05 were considered to have significant activity.

### Pathway enrichment analysis

Enricher function from clusterprofiler^72^ was used to perform over representation analysis (ORA) on genes (based on EntrezID) shared by *in vitro* myCAF-driven PDAC markers and *in vivo* myCAF-dependent PDAC markers or *in vitro* myCAF markers and *in vivo* myCAF markers. Gene lists are included in **Supplementary Tables 2-3**.

### Single-cell RNA-sequencing of murine PDAC FTC and FT tumours

scRNA-seq data of murine PDAC tumours are available at the GEO under the accession number GSE342740. scRNA-seq analyses are included in **Supplementary Table 1**. CellRanger pipeline (10x Genomics) was used to align raw fastq files to Mouse mm10 (2020-A) reference genome amended with the reporter genes: *Tdtomato* and *Cre*. For ambient RNA removal the *raw_feature_bc_matrix* files were used as input for CellBender^77^ with fpr=0.01. Each sample underwent standard quality control using Scanpy^78^. For preliminary quality control, cells were filtered based on total counts (> 60000 counts), gene counts (<100 genes), and mitochondrial gene content (>5%). SOLO model^79^, implemented in scvi-tools, was used to estimate doublets. After doublet removal, all samples were concatenated and integrated using Harmony^80^ and “SampleID” was used as a batch key. After integration, data was clustered using Leiden algorithm^81^ based on top 4000 highly variable genes and top 30 PCs. Cell clusters were annotated based on known gene marker and reporter gene expression. To identify malignant cells, a python implementation of inferCNV of the Trinity CTAT Project (https://github.com/broadinstitute/inferCNV) was used to estimate the copy number status in each cell type with fibroblasts as reference key and a 250-genes window size. DEA was applied using the Model-based Analysis of Single-cell Transcriptomics (MAST) method^82^, were “SampleID” used as a latent variable. Genes for GSEA were ranked based on the log2 fold change.

### Pseudotime trajectory analysis

Palantir^83^ was used with default parameters to estimate the potential cell trajectories of CAFs in the FTC samples. A starting cell for the trajectory was identified by choosing a cell at the maximal point in the myCAF subcluster, while the terminal cells were automatically identified by Palantir.

### Single-cell RNA-sequencing of murine PDAC tumours in C57BL/6J mice

scRNA-seq data of murine PDAC tumours (n=8 moPDAC-young only) in C57BL/6J mice were previously published^84^ and are available at the GEO under the accession number GSE263595. Raw data were pre-processed and analysed using the same pipeline and parameters mentioned in the scRNA-seq data analysis section.

### RNA-sequencing of CAF-locked lines cultured in 2D

RNA-seq data of iCAF-locked and myCAF-locked PSCs cultured in 2D mice are available at the GEO under the accession number GSEA342726^27^. Data were analysed using the same steps and parameters mentioned in the bulk RNA-seq data analysis section.

### Hypergeometric test and overrepresenting analysis

Hypergeometric test, odds ratio and Jaccard index were calculated for overlapping genes between the selected pairs of gene lists using GeneOverlap R package^85^. As done previously^32^, the universe was set to 18,000 to represent the number of curated protein-coding genes and including *CDKN2A* and *CDKN2B* (https://bioconductor.org/packages/release/bioc/html/GeneOverlap.html). Over representation analysis (ORA) was run using enricher function from clusterprofiler R package on the overlapping genes.

### Statistical analysis

GraphPad Prism software, customized R scripts and Jupyter notebooks were used for graphical representation of data. Statistical analysis was performed using non-parametric Mann-Whitney test, non-parametric Kruskal-Wallis adjusted for multiple comparisons, unpaired two-sided Welch’s t-test, bootstrap test, hypergeometric test and Fisher’s exact test. Adjustment for multiple comparisons was applied where relevant. All statistical details of experiments, and how significance was defined, are specified in the figure legends and/or panel figures, including the number of replicates.

## ACKNOWLEDGEMENTS

The authors would like to thank the CRUK-CI BRU, Genomics, Bioinformatics, Flow Cytometry, Pre-genome editing, and Histology core facilities. The authors would also like to acknowledge Professor Fearon (previously at CRUK-CI) for sharing the *Fap*-tTA mouse line, Professor Winton (previously at CRUK-CI) for sharing the *Rosa26*-tdTomato mouse line, Professor Linterman (Babraham Institute and Malaghan Institute of Medical Research) for sharing the tet-O-Cre-LC1 mouse line, and Professor Denton (Imperial College London) for advice on breeding strategy and dox regimen. This work was mainly supported by G.B. Cancer Research UK institutional grant (A27463), which also supported J.S.M. and S.M. This work was also supported by a UKRI Future Leaders Fellowship of which GB is recipient and that also supported W.L. and E.G.L. The Pancreatic Cancer Research Fund and the US Department of Defense (PCARP grant) supported G.M. and J.S.M. A NCI-CRUK Cancer Grand Challenge grant supported M.J., W.K.L. and S.H. A Pancreatic Cancer UK Future Leaders Academy grant supported P.S.W.C. J.A.H. was supported by a Harding Distinguished Postgraduate Programme PhD studentship (Cambridge Trust).

## AUTHOR CONTRIBUTIONS

E.G.L and P.S.W.C. designed the experiments, conducted the experiments, and revised the paper. M.J., M.Z., K.B., W.L., J.S.M., W.K.L, S.H., J.A.H., S.M., D.M., G.M., P.M.J. and A.A.J. conducted the experiments. M.V. oversaw the mouse studies. M.D. advised on rheology experiments. P.D.W.K co-supervised M.J. and advised on statistical analyses. G.B conceptualized and supervised the study, designed the experiments, conducted the experiments, and wrote the paper.

## DECLARATION OF INTERESTS

No competing interests.

## SUPPLEMENTARY FIGURE LEGENDS AND SUPPLEMENTARY TABLES

**Supplementary Figure 1.**
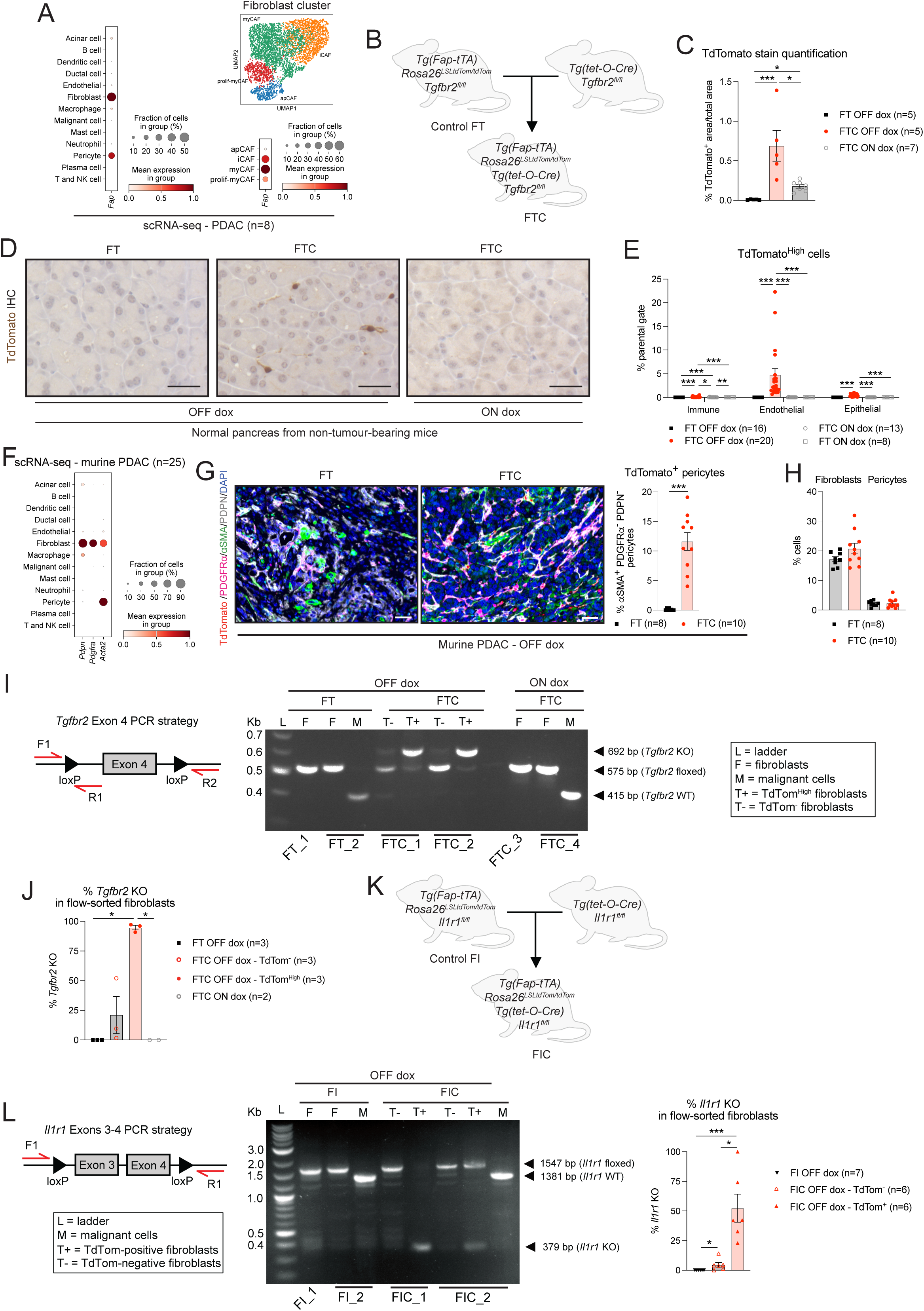
Characterisation of novel mouse models for depletion of *Fap*-expressing TGF-β- or IL-1-dependent fibroblasts in PDAC. **(A)** Dot plots of the scaled average expression of *Fap* in all cell type (left) or cancer-associated fibroblast (CAF, right) clusters from pancreatic ductal adenocarcinoma (PDAC) tumours (n=8) analysed by single-cell RNA-sequencing (scRNA-seq). The colour intensity represents the expression level, and the size of the dots represents the percentage of expressing cells. A Uniform Manifold Approximation and Projection (UMAP) plot of the fibroblast cluster is provided for reference. The scRNA-seq data is from Araos Henríquez et al. **(B)** Schematic summarising the breeding strategy to obtain *Fap-tTa; Rosa26^LSL-tdTom/tdTom^*; *Tgfbr2^fl/fl^* (hereafter, FT) and *Fap-tTa; tet-O-Cre; Rosa26^LSL-tdTom/tdTom^; Tgfbr2^fl/fl^* (hereafter, FTC) mice. **(C)** Quantification of TdTomato stains in FT (n=5) and FTC (n=5) normal pancreata 6-weeks after doxycycline (dox) withdrawal and in normal pancreata of FTC mice on dox (n=7). Results show mean ± SEM. * *P adj* < 0.05; *** *P adj* < 0.001, Kruskal-Wallis test. **(D)** Representative TdTomato stains in FT and FTC normal pancreata 6-weeks after dox withdrawal and in normal pancreata of FTC mice on dox. Scale bars, 100 μm. **(E)** Flow cytometric analyses of TdTomato^High^ immune (PDPN^-^CD31^-^CD45^+^), endothelial (PDPN^-^CD45^-^CD31^+^) and epithelial (PDPN^-^CD31^-^CD45^-^ EpCAM^+^) cells from live singlets in FT (n=16) and FTC (n=20) PDAC after dox withdrawal or in FT (n=8) and FTC (n=13) PDAC on dox. Results show mean ± SEM. * *P adj* < 0.05; ** *P adj* < 0.01; *** *P adj* < 0.001, Kruskal-Wallis test. **(F)** Dot plot of the scaled average expression of *Pdpn, Pdgfra and Acta2* (encoding alpha smooth muscle actin, αSMA) in all cell type from PDAC tumours (n=8) analysed by scRNA-seq. The colour intensity represents the expression level, and the size of the dots represents the percentage of expressing cells. The scRNA-seq data is from Araos Henríquez et al. **(G)** Left: Representative multiplex immunofluorescence (mIF) images of TdTomato (red), podoplanin (PDPN, white), αSMA (green), platelet-derived growth factor receptor alpha (PDGFRα, pink) and DAPI (nuclear stain, blue) in FT and FTC PDAC tissues after dox withdrawal. Scale bars, 50 μm. Right: Quantification of TdTomato^+^αSMA^+^PDGFRα^-^PDPN^-^ECAD^-^DAPI^+^ cells relative to αSMA^+^ECAD^-^DAPI^+^ cells in FT (n=8) and FTC (n=10) PDAC tissues after dox withdrawal. Results show mean ± SEM. *** *P* < 0.001, Mann-Whitney test. **(H)** Quantification of DAPI^+^ fibroblasts (PDPN^+^ECAD^-^) and DAPI^+^ pericytes (αSMA^+^PDGFRα^-^PDPN^-^ECAD^-^) relative to all (DAPI^+^) cells in FT (n=8) and FTC (n=10) PDAC tissues after dox withdrawal. Results show mean ± SEM. **(I)** Left: Schematic of PCR strategy to validate *Tgfbr2* knock out (KO) in FTC models. Right: Representative PCR results of *Tgfbr2* allele status from TdTomato^High^ (T+) and TdTomato^-^ (T-) fibroblasts flow-sorted from FTC PDAC after dox withdrawal, or fibroblasts (F) and malignant cells (M) flow-sorted from FT PDAC after dox withdrawal and from FTC PDAC on dox. *Tgfbr2* wild type (WT) allele at 415 bp, floxed allele at 575 bp, KO allele at 692 bp. **(J)** PCR-based quantification of *Tgfbr2* KO percentage in TdTomato-high (TdTom^High^) and TdTomato^-^ (TdTom^-^) fibroblasts flow-sorted from FTC PDAC after dox withdrawal (n=3), as well as fibroblasts flow-sorted from FT after dox withdrawal (n=3) and FTC PDAC on dox (n=2). * *P adj* < 0.05, Kruskal-Wallis test. **(K)** Schematic summarising the breeding strategy to obtain *Fap-tTa; Rosa26^LSL-tdTom/tdTom^*; *Il1r1^fl/fl^* (hereafter, FI) and *Fap-tTa; tet-O-Cre; Rosa26^LSL-tdTom/tdTom^*; *Il1r1^fl/fl^* (hereafter, FIC) mice. **(L)** Left: Schematic of PCR strategy to validate *Il1r1* KO in FIC models. Middle: Representative PCR results of *Il1r1* allele status from TdTomato^+^ (T+), TdTomato^-^ (T-) and malignant cells (M) flow-sorted from FIC PDAC after dox withdrawal, or fibroblasts (F) and malignant cells (M) flow-sorted from FI PDAC after dox withdrawal. *Il1r1* WT allele at 1381 bp, floxed allele at 1547 bp, KO allele at 379 bp. Right: PCR-based quantification of *Il1r1* KO percentage in TdTomato^+^ (TdTom^+^) and TdTomato^-^ (TdTom^-^) fibroblasts flow-sorted from FIC PDAC after dox withdrawal (n=6), as well as fibroblasts flow-sorted from FI after dox withdrawal (n=7). * *P adj* < 0.05, *** *P adj* < 0.001; Kruskal-Wallis test.

**Supplementary Figure 2.**
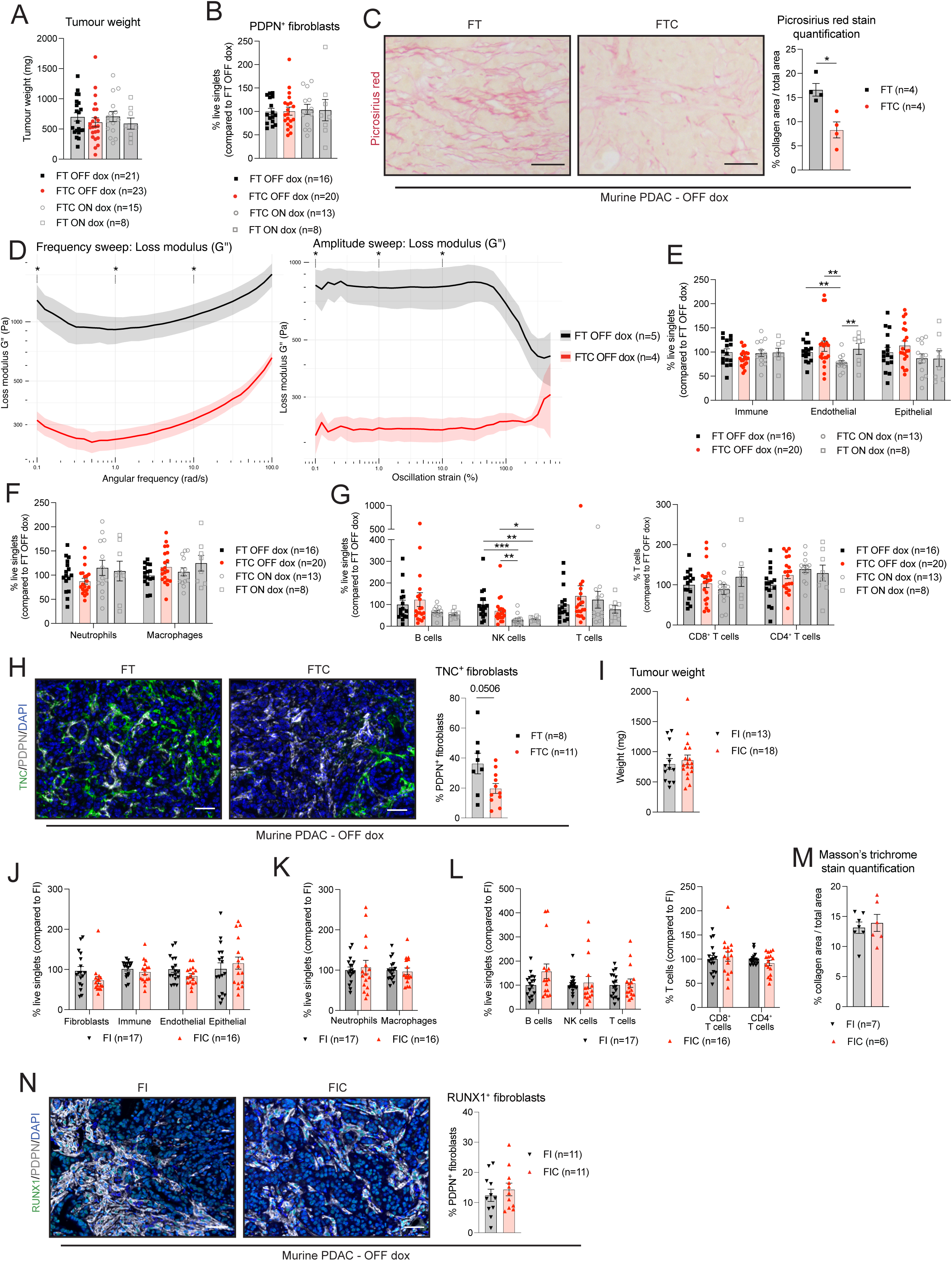
Disruption of TGF-β or IL-1 signalling in *Fap*-expressing fibroblasts differentially remodels PDAC tumours. **(A)** Weights measured at experimental endpoint of FT (n=21, 40.2 ± 2.8 days post-transplant) and FTC (n=23, 39.6 ± 3.4 days post-transplant) PDAC after dox withdrawal or of FT (n=8, 37.5 ± 3.6 days post-transplant) and FTC (n=15, 41.9 ± 5.8 days post-transplant) PDAC on dox. Results show mean ± SEM from 8 separate experiments. No statistical difference was found, as calculated by Kruskal-Wallis test. **(B)** Flow cytometric analysis of PDPN^+^ fibroblasts (CD45^-^CD31^-^EpCAM^-^) from live singlets in FT (n=16) and FTC (n=20) PDAC after dox withdrawal or in FT (n=8) and FTC (n=13) PDAC on dox. Results show mean ± SEM. No statistical difference was found, as calculated by Kruskal-Wallis test. **(C)** Left: Representative picrosirius red stains in FT and FTC PDAC after dox withdrawal. Scale bars, 100 μm. Right: Picrosirius red stain quantification in FT (n=4) and FTC (n=4) PDAC after dox withdrawal. Results show mean ± SEM. * *P* < 0.05; Mann-Whitney test. **(D)** Oscillatory rheological properties of *ex vivo* FT (n=5) and FTC (n=4) PDAC tumours after dox withdrawal under hydrated conditions at 37 °C: loss moduli from frequency (left) and amplitude (right) sweeps. Amplitude sweeps were performed at a constant angular frequency of 1 rad/s, while frequency sweeps were performed at a constant strain of 0.5%. * *P* < 0.05, unpaired two-sided Welch’s *t*-test with 0.1%, 1% and 10% strain for amplitude sweeps at 0.1, 1, and 10 rad/s for frequency sweeps. **(E)** Flow cytometric analysis of immune cells (CD45^+^CD31^-^), endothelial cells (CD31^+^CD45) and epithelial cells (CD45^-^CD31^-^PDPN^-^EpCAM^+^) from live singlets in FT (n=16) and FTC (n=20) PDAC after dox withdrawal or in FT (n=8) and FTC (n=13) PDAC on dox. Results show mean ± SEM. ** *P adj* < 0.01; Kruskal-Wallis test. **(F)** Flow cytometric analysis of neutrophils (CD45^+^CD11b^+^Gr1^+^) and macrophages (CD45^+^CD11b^+^Gr1^-^F4/80^+^) from live singlets in FT (n=16) and FTC (n=20) PDAC after dox withdrawal or in FT (n=8) and FTC (n=13) PDAC on dox. Results show mean ± SEM. No statistical difference was found, as calculated by Kruskal-Wallis test. **(G)** Flow cytometric analysis of T cells (CD45^+^CD3ε^+^TCRβ^+^), NK cells (CD45^+^NK1.1^+^) and B cells (CD45^+^CD19^+^) from live singlets (left) and CD4^+^ and CD8^+^ T cells from the CD45^+^CD3ε^+^TCRβ^+^ parental gate (right) in FT (n=16) and FTC (n=20) PDAC after dox withdrawal or in FT (n=8) and FTC (n=13) PDAC on dox. Results show mean ± SEM. * *P adj* < 0.05, ** *P adj* < 0.01, *** *P adj* < 0.001; Kruskal-Wallis test. **(H)** Left: Representative mIF images of TNC (green), PDPN (white) and DAPI (blue) in FT and FTC PDAC tissues after dox withdrawal. Scale bars, 50 μm. Right: Quantification of TNC^+^PDPN^+^CK19^-^DAPI^+^ fibroblasts relative to CK19^-^PDPN^+^DAPI^+^ fibroblasts in FT (n=8) and FTC (n=11) PDAC tissues after dox withdrawal. Results show mean ± SEM. Mann-Whitney test. **(I)** Weights measured at experimental endpoint of FI (n=13, 43.4 ± 2.0 days post-transplant) and FIC (n=18, 44.6 ± 4.5 days post-transplant) PDAC after dox withdrawal. Results show mean ± SEM. No statistical difference was found, as calculated by Mann-Whitney test. **(J)** Flow cytometric analysis of fibroblasts, immune cells, endothelial cells and epithelial cells from live singlets in FI (n=17) and FIC (n=16) PDAC after dox withdrawal. Results show mean ± SEM. No statistical difference was found, as calculated by Mann-Whitney test. **(K)** Flow cytometric analysis of neutrophils and macrophages from live singlets in FI (n=17) and FIC (n=16) PDAC after dox withdrawal. Results show mean ± SEM. No statistical difference was found, as calculated by Mann-Whitney test. **(L)** Flow cytometric analysis of T cells, NK cells and B cells from live singlets (left) and CD4^+^ and CD8^+^ T cells from the parental gate (right) in FI (n=17) and FIC (n=16) PDAC after dox withdrawal. Results show mean ± SEM. No statistical difference was found, as calculated by Mann-Whitney test. **(M)** Quantification of Masson’s trichrome stain in FI (n=7) and FIC (n=6) PDAC after dox withdrawal. Results show mean ± SEM. No statistical difference was found, as calculated by Mann-Whitney test. **(N)** Left: Representative mIF images of RUNX1 (green), PDPN (white) and DAPI (blue) in FI and FIC PDAC tissues after dox withdrawal. Scale bars, 50 μm. Right: Quantification of RUNX1^+^PDPN^+^CK19^-^DAPI^+^ fibroblasts relative to PDPN^+^CK19^-^DAPI^+^ fibroblasts in FI (n=11) and FIC (n=11) PDAC tissues after dox withdrawal. Results show mean ± SEM. No statistical difference was found, as calculated by Mann-Whitney test.

**Supplementary Figure 3.**
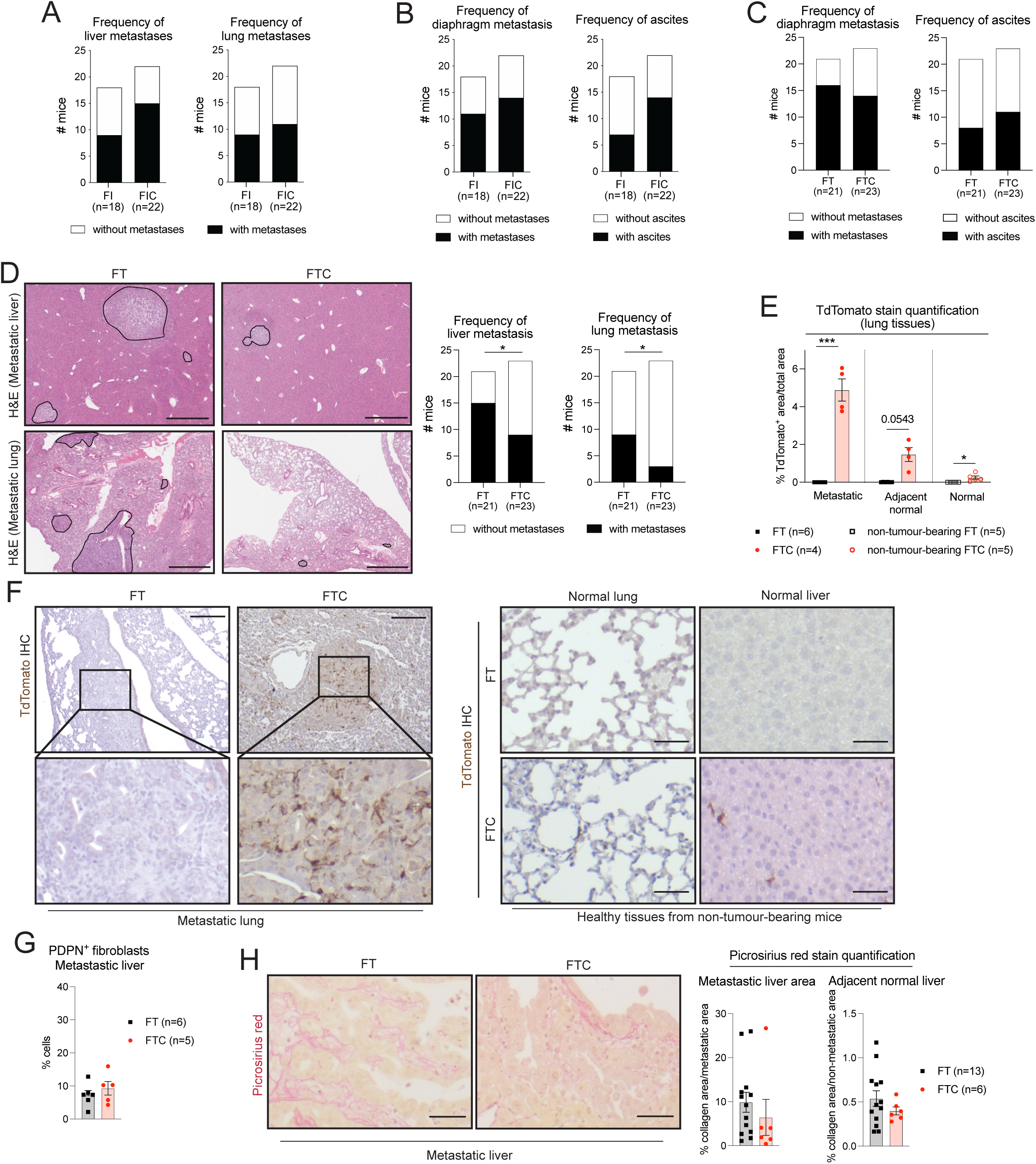
Depletion of myCAFs impairs PDAC liver and lung metastasis formation. (A-B) Number of FI (n=18) and FIC (n=22) mice after dox withdrawal with or without metastases in the liver and lungs **(A)** or with or without metastases in the diaphragm and with or without ascites **(B)**. No statistical difference was found, as calculated by Fisher’s exact test. **(C)** Number of FT (n=21) and FTC (n=23) mice after dox withdrawal with or without metastases in the diaphragm and with or without ascites. No statistical difference was observed, as calculated by Fisher’s exact test. **(D)** Left: Representative Haematoxylin and Eosin (H&E) stains of lung and liver metastatic tissues in tumour-bearing FT and FTC mice after dox withdrawal. Scale bars, 900 μm and 1 mm for liver and lung, respectively. Right: Number of FT (n=21) and FTC (n=23) mice after dox withdrawal with or without metastases in the liver and lungs. * *P* < 0.05, Fisher’s exact test. **(E)** TdTomato stain quantification in lung metastatic and adjacent normal tissue from FT (n=6) and FTC (n=4) tumour-bearing mice, and in normal lung tissue in non-tumour-bearing FT (n=5) and FTC (n=5) mice, after dox withdrawal. Results show mean ± SEM. * *P adj* < 0.05; *** *P adj* < 0.001; Kruskal-Wallis test. **(F)** Representative TdTomato IHC stains of lung metastatic tissues in tumour-bearing FT and FTC mice (left) or of lung and liver normal tissues in non-tumour-bearing FT and FTC mice (right) after dox withdrawal. Scale bars, 100 μm. Inserts, magnifications. **(G)** Quantification of PDPN^+^ fibroblasts relative to all (DAPI^+^) cells in FT (n=6) and FTC (n=5) liver metastatic tissues after dox withdrawal. Results show mean ± SEM. No statistical difference was found, as calculated by Mann-Whitney test. **(H)** Left: Representative picrosirius red stains in FT and FTC PDAC tissues after dox withdrawal. Scale bars, 25 μm. Middle/Right: Picrosirius red stain quantification in liver metastatic tissues (middle) and in adjacent normal liver tissues from FT (n=6) and FTC (n=5) tumour-bearing mice after dox withdrawal. Results show mean ± SEM. No statistical difference was observed, as calculated by Mann-Whitney test.

**Supplementary Figure 4.**
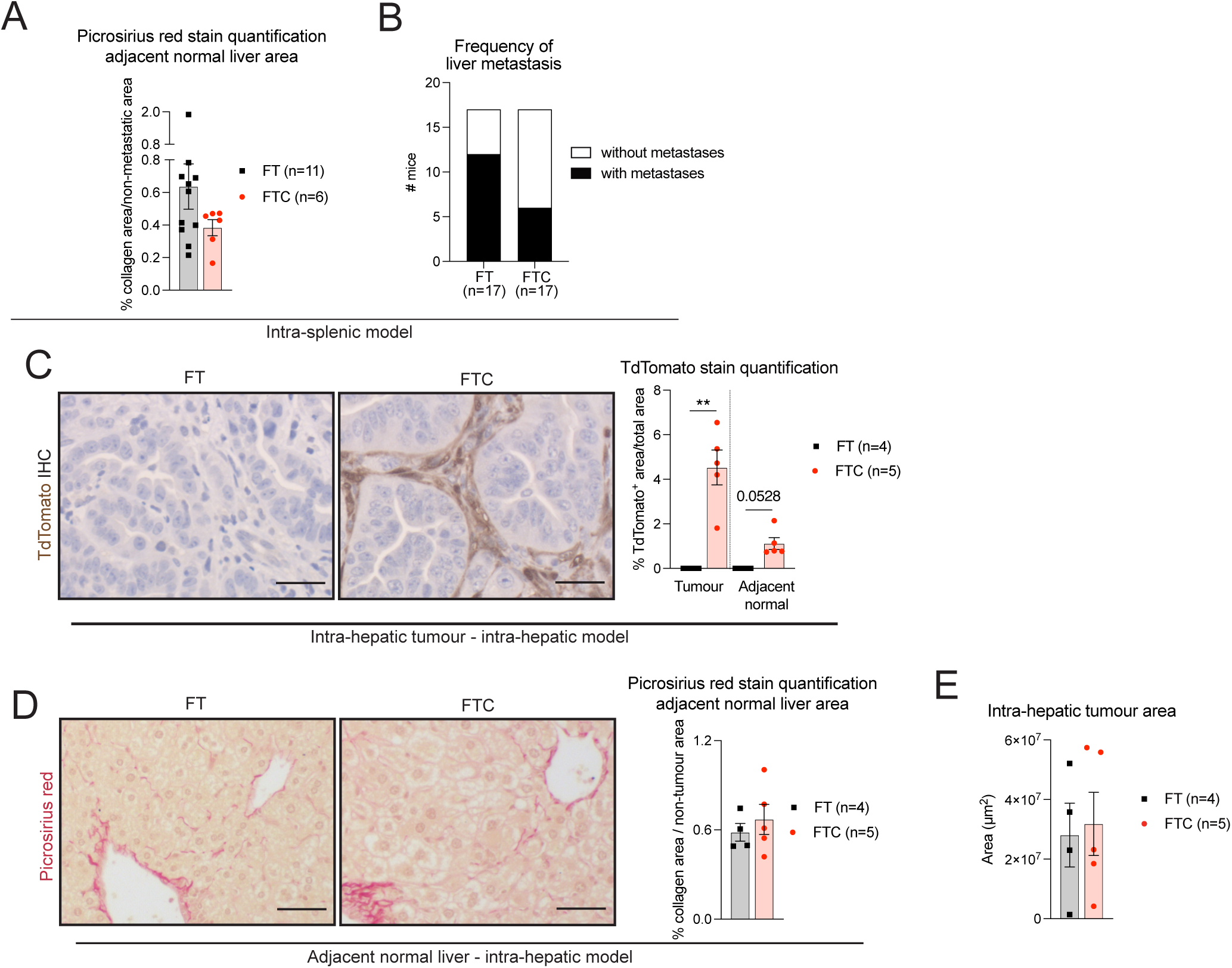
Disruption of TGF-β signalling in *Fap*-expressing fibroblasts does not impair PDAC liver metastatic outgrowth. **(A)** Picrosirius red stain quantification in adjacent normal liver tissues from intra-splenic FT (n=11) and FTC (n=6) mice after dox withdrawal. Results show mean ± SEM. No statistical difference was found, as calculated by Mann-Whitney test. **(B)** Number of intra-splenic FT (n=17) and FTC (n=17) mice after dox withdrawal with or without metastases in the liver. No statistical difference was observed, as calculated by Fisher’s exact test. **(C)** Left: Representative TdTomato IHC stains of FT and FTC intra-hepatic tumours after dox withdrawal. Scale bars, 25 μm. Right: TdTomato stain quantification in intra-hepatic tumours from intra-hepatic FT (n=4) and FTC (n=5) models after dox withdrawal. Results show mean ± SEM. ** *P adj* < 0.01, Kruskal-Wallis test. **(D)** Left: Representative picrosirius red stains of adjacent normal liver tissues from FT and FTC intra-hepatic models after dox withdrawal. Scale bars, 25 μm. Right: Picrosirius red stain quantification in adjacent normal liver tissues from intra-hepatic FT (n=4) and FTC (n=5) models after dox withdrawal. Results show mean ± SEM. No statistical difference was found, as calculated by Mann-Whitney test. **(E)** Areas measured at experimental endpoint of FT (n=4, 33 ± 0 days post-transplant) and FTC (n=5, 32.6 ± 0.9 days post-transplant) intra-hepatic tumours after dox withdrawal. Results show mean ± SEM. No statistical difference was found, as calculated by Mann-Whitney test.

**Supplementary Figure 5.**
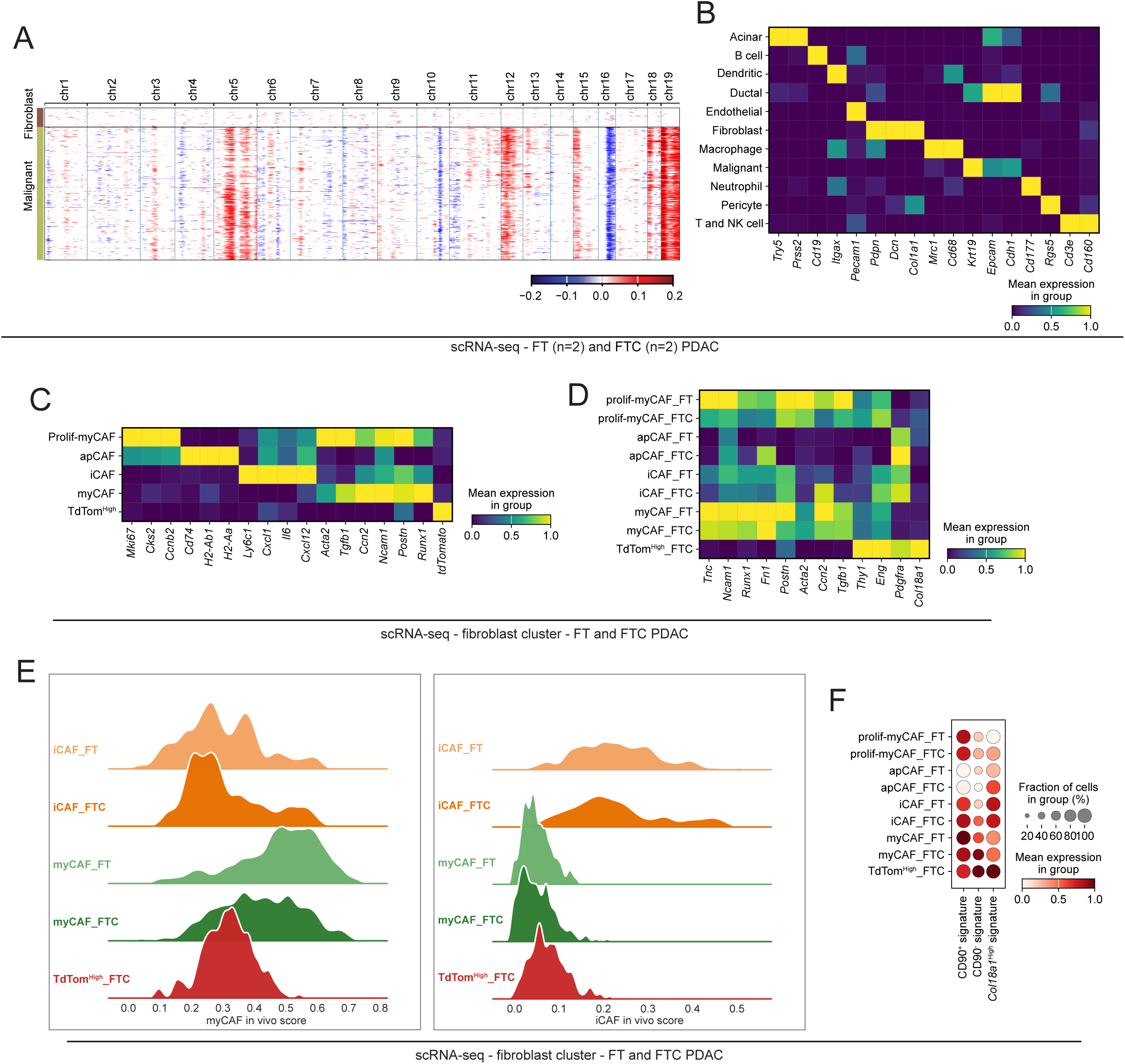
Depletion of myCAFs drives broad transcriptional changes in the primary TME. **(A)** Heatmap showing large-scale copy number variation (CNV) profile of the fibroblast and epithelial malignant cell clusters identified by scRNA-seq of murine FT (n=2) and FTC (n=2) PDAC. The colour coding represents the CNV level based on a sliding window of 250 gene expression. Amplifications are shown in red, and deletions are shown in blue. The fibroblast cluster was used as reference cell cluster. **(B)** Heatmap of scaled expression of cell type-specific markers in each cell cluster of in FT and FTC PDAC, as analysed by scRNA-seq. Data are scaled such that the cluster with the lowest average expression = 0 and the highest = 1 for each gene. **(C)** Heatmap of scaled expression of CAF cluster-defining markers in each CAF cluster of FT and FTC PDAC, as analysed by scRNA-seq. Data are scaled such that the cluster with the lowest average expression = 0 and the highest = 1 for each gene. **(D)** Heatmap of scaled expression of CAF markers in each CAF cluster of FT and FTC PDAC, as analysed by scRNA-seq. Data are scaled such that the cluster with the lowest average expression = 0 and the highest = 1 for each gene. **(E)** Ridge plots showing the distribution of *in vivo* myCAF (left) and *in vivo* iCAF (right) gene signature AUCell scores across selected CAF clusters from scRNA-seq datasets of FT and FTC PDAC tumours. Shaded areas represent the density distribution of AUCell scores within each CAF subtype. **(F)** Dot plot of scaled expression of murine i*n vivo* CD90^+^ myCAF, CD90^-^ myCAF and *Col18a1*^High^ CAF signatures in the fibroblast cluster of FT and FTC PDAC. The colour intensity represents the expression level, and the size of the dots represents the percentage of expressing cells. The CD90^+^ and CD90^-^ myCAF signatures are from Mucciolo et al. The *Col18a1*^High^ CAF signature is from Kuhn et al.

**Supplementary Figure 6.**
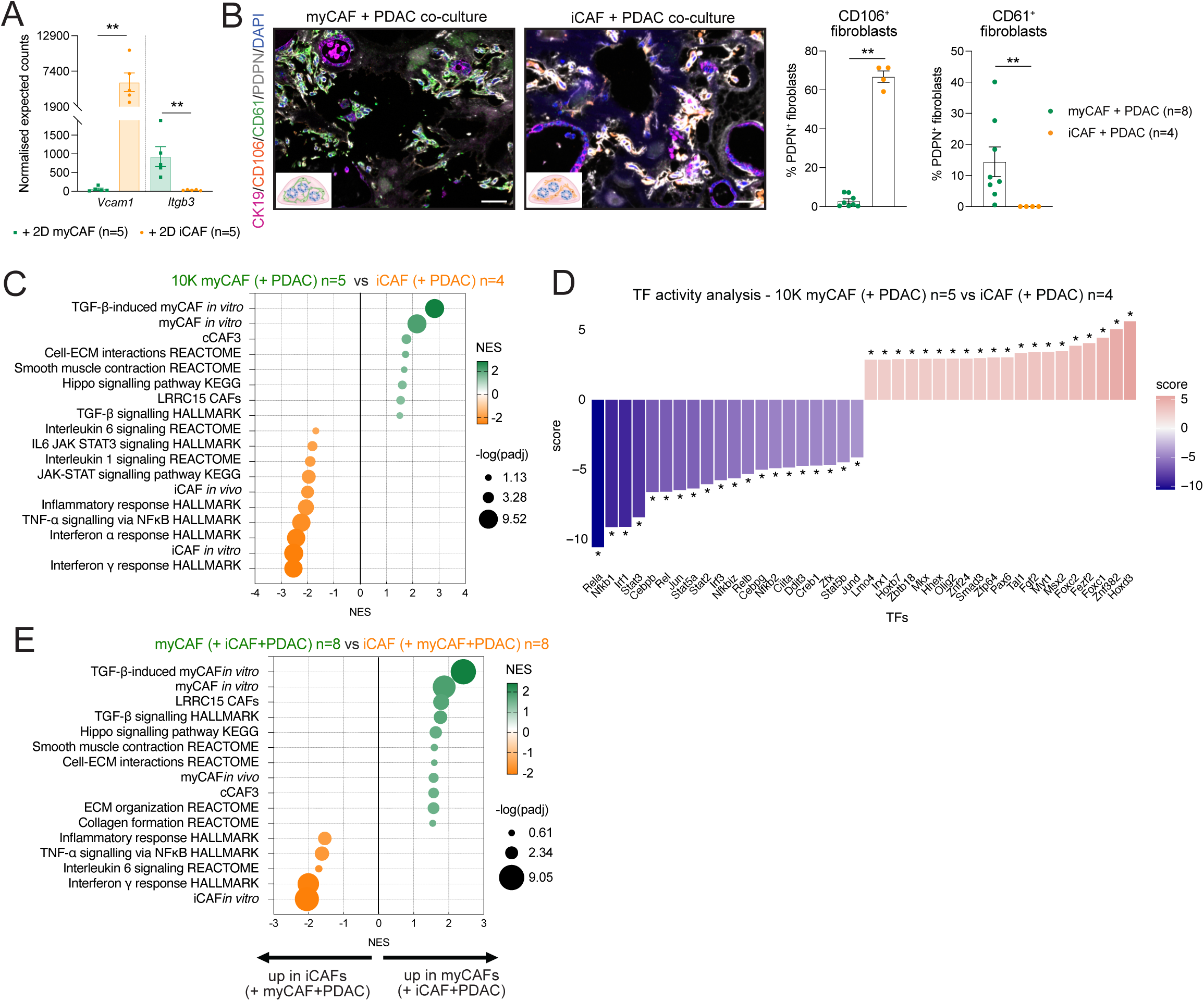
myCAF-locked PSCs capture TGF-β-dependent CAF signalling observed *in vivo*. **(A)** RNA-sequencing (RNA-seq) expression of *Vcam1* (encoding CD106) and *Itgb3* (encoding CD61) in myCAF-locked PSCs (myCAF, n=5) and iCAF-locked PSCs (iCAF, n=5) cultured in 2D. Results show mean ± SEM. ** *P* < 0.01, Mann-Whitney test. RNA-seq data are from Pelicano et al. **(B)** Left: Representative mIF images of CK19 (pink), PDPN (white), CD106 (orange), CD61 (green) and DAPI (blue) stains of PDAC organoid/myCAF co-cultures or PDAC organoid/iCAF co-cultures after 4 days in culture. Scale bars = 50 μm. Right: quantification of CD106^+^CD61^-^PDPN^+^CK19^-^DAPI^+^ and CD61^+^CD106^-^PDPN^+^CK19^-^DAPI^+^ cells over PDPN^+^CK19^-^DAPI^+^ cells in PDAC organoid/myCAF co-cultures (n=8) or PDAC organoid/iCAF co-cultures (n=4). Results show mean ± SEM. ** *P* < 0.01, Mann-Whitney test. **(C)** Selected significantly upregulated and downregulated pathways identified by Gene Set Enrichment Analysis (GSEA) of myCAFs co-cultured with PDAC organoids (n=5) compared to iCAFs co-cultured with PDAC organoids (n=4). The *in vivo* iCAF signature is from Elyada et al. The *in vitro* iCAF and myCAF signatures are from Öhlund et al. The TGF-β-induced myCAF *in vitro* signature is from Mucciolo and Araos Henríquez et al. The LRRC15^+^ CAF signature was obtained is from Dominguez et al. The cCAF3 signature is from McAndrews et al. In this experiment myCAFs were plated at half density respective to iCAFs, noted by the 10K. **(D)** Activity score of the top 20 most activated and most deactivated transcription factors (TFs) in myCAFs co-cultured with PDAC organoids (n=5) compared to iCAFs co-cultured with PDAC organoids (n=4). * *P* < 0.05, univariate linear model (ulm) method. In this experiment myCAFs were plated at half density respective to iCAFs, noted by the 10K. **(E)** Selected significantly upregulated and downregulated pathways identified by GSEA of myCAFs compared to iCAFs both flow-sorted from triple co-cultures of PDAC organoids with iCAFs and myCAFs (n=8). The *in vivo* myCAF signature is from Elyada et al.

**Supplementary Figure 7.**
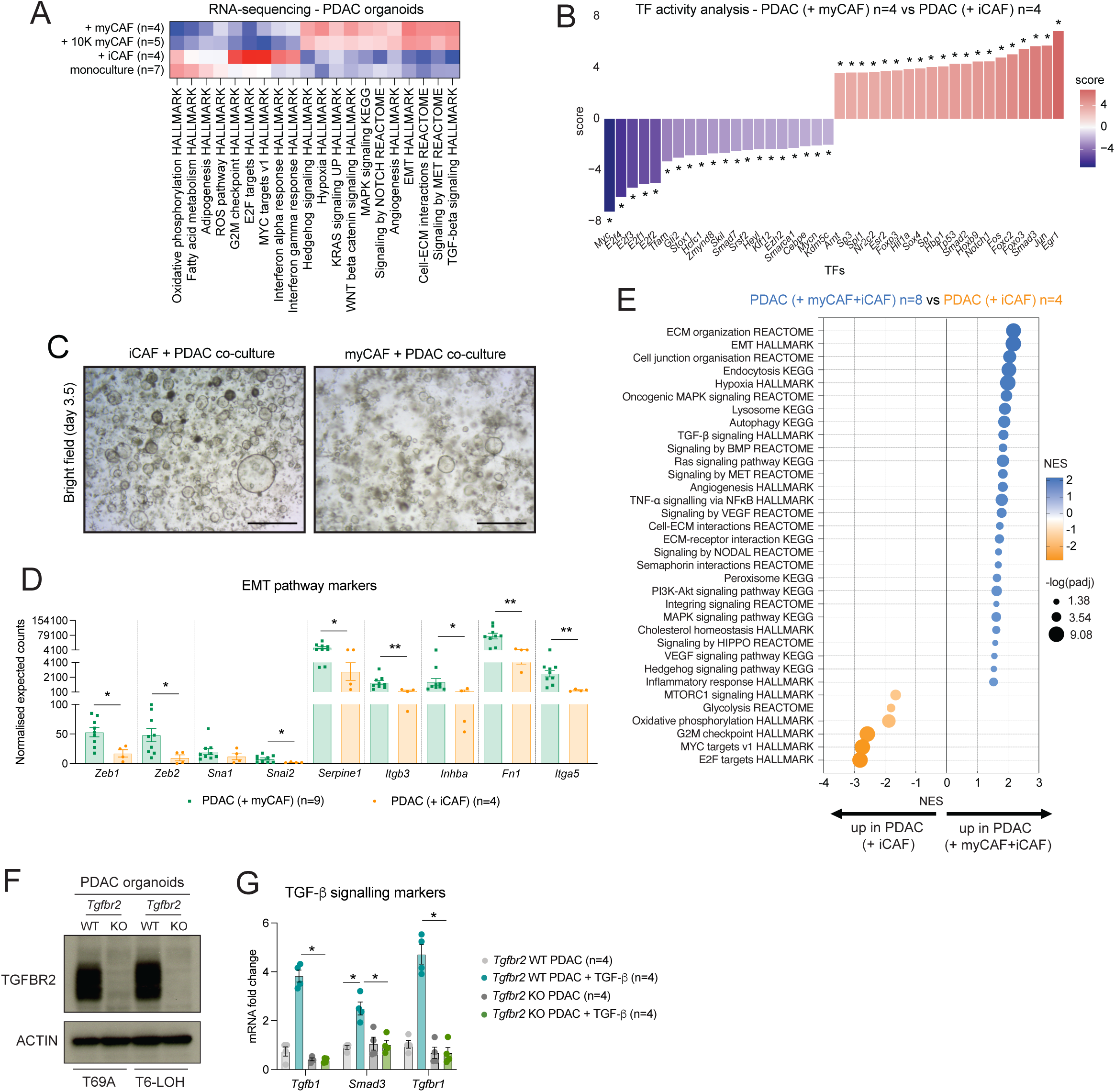
myCAF-locked PSCs upregulate EMT signalling in PDAC malignant cells. **(A)** Heatmap showing Gene Set Variation Analysis (GSVA) enrichment scores (averaged per group) of selected pathways in PDAC organoids in monoculture (n=7), in co-culture with iCAFs (n=4), in co-culture with myCAFs (n=4) and in co-culture with 10K myCAFs (n=5), as assessed by RNA-seq. **(B)** Activity score of the top 20 most activated and most deactivated TFs in PDAC organoids co-cultured with myCAFs (n=4) versus PDAC organoids co-cultured with iCAFs (n=4). * *P* < 0.05, ulm method. **(C)** Bright field images of PDAC organoids in co-culture with iCAFs (left) or myCAFs (right) after 3.5 days in culture. Scale bars, 300 μm. For both CAF-locked lines, initial plating was 20K/Matrigel dome. **(D)** RNA-seq expression of epithelial-to-mesenchymal transition (EMT) markers in PDAC organoids co-cultured with myCAFs (n=9, with n=5 co-cultured with 10K myCAFs and n=4 co-cultured with 20K myCAFs) or iCAFs (n=4). Results show mean ± SEM. * *P* < 0.05; ** *P* < 0.01, Mann-Whitney test. **(E)** Selected significantly upregulated and downregulated pathways identified by GSEA of PDAC organoids co-cultured with myCAFs and iCAFs (PDAC triple cocx, n=8) compared to PDAC organoids co-cultured with iCAFs (PDAC + iCAFs, n=4). **(F)** Western blot analysis of the TGF-β receptor 2 (TGFBR2) in *Tgfbr2* WT and *Tgfbr2* KO PDAC organoids cultured for 3 days. ACTIN, loading control. **(G)** qPCR analysis of TGF-β signalling mediators *Tgfb1, Smad3* and *Tgfbr1* in *Tgfbr2* WT and *Tgfbr2* KO PDAC organoids cultured for 3 days with or without 2 ng/mL recombinant TGF-β (n=4/condition). Results show mean ± SEM. * *P adj* < 0.05, Kruskal-Wallis test.

### LIST OF SUPPLEMENTARY TABLES

**Table S1.** Single cell RNA-sequencing of murine PDAC from FTC and FT mice.

**Table S2.** RNA-sequencing of flow-sorted CAF-locked PSCs from double and triple co-cultures with PDAC organoids.

**Table S3.** RNA-sequencing of flow-sorted organoids from double and triple co-cultures with CAF-locked PSCs and from monocultures.

## Notes

### Competing Interest Statement

The authors have declared no competing interest.

